# Rapid speciation in Harlequin Toads (*Anura*: *Bufonidae*) endemic to the Sierra Nevada de Santa Marta

**DOI:** 10.64898/2026.05.28.728521

**Authors:** Juan Ramirez-Romero, Laura M. Eslava, Fabian C. Salgado-Roa, José D. Barros-Castañeda, Lucas S. Barrientos, Andrew J. Crawford, Carolina Pardo-Díaz, Luis Alberto Rueda-Solano, Camilo Salazar

## Abstract

The Neotropical genus *Atelopus* has experienced a drastic population decline in recent decades. Despite this, a knowledge gap remains regarding the conservation genetic status, phylogeography, and demographic history of most of its species, especially those endemic to areas with limited access like the Sierra Nevada de Santa Marta in Colombia. In this genomic study, we inferred phylogenetic relationships, demographic history, and gene flow among four endemic *Atelopus* morphospecies in the Sierra Nevada de Santa Marta-SNSM. Additionally, we compared the effective population size (N_e_) estimates with the available population census data. NextRAD was used to obtain genomic data from 95 individuals collected at five sites in the SNSM and two in the Colombian Pacific (outgroup). The morphospecies recently diverged in a scenario without gene flow and were recovered as monophyletic. Their phylogenetic relationships were discordant, which is attributed to the presence of incomplete lineage sorting-ILS, which would also explain their shared ancestry among them. The lack of gene flow as well as the recent divergence times given by demography suggests a recent and rapid speciation. However, the reproductive isolation mechanisms that promote or maintain the species boundaries in this group remain unknown and require further investigation. We suggest that this process may have been influenced by the complex topography of the SNSM, traits such as high philopatry, low dispersal ability, and behavioral factors such as habitat preference or to factors related to genetic architecture that influenced the rapid formation of reproductive barriers among populations. Additionally, a pattern of population decline was observed around 200.000 years ago, with recent increases in three morphospecies. Despite the reduction in effective population size, no signs of inbreeding were detected. However, for *A. laetissimus*, the only species surveyed, the estimated value of N_e_ and its implications should be interpreted with caution. Ultimately, our findings reveal an evolutionary history shaped by a burst of diversification and abrupt reproductive isolation, highlighting how the resulting endemism and restricted genetic connectivity shape the unique evolutionary trajectory and vulnerability of this threatened montane species.

**Significance Statement:** The mechanisms driving rapid speciation in montane ecosystems remain a central question in evolutionary biology. This study provides crucial genomic insights into the diversification of four endemic *Atelopus* species in an isolated Neotropical massif. We reveal a compelling evolutionary scenario where species diverge rapidly in absence of gene flow. This rapid speciation was likely facilitated by complex topography, environmental heterogeneity, and ecological and behavioral differences among species. Nevertheless, the evolutionary processes that limited gene flow among SNSM species have not yet been identified.

## Introduction

Changes in the geographic and ecological landscape can shape species distribution and diversity, as well as their demographic and evolutionary history (Gascuel et al., 2015; Muñoz-Ortiz et al., 2015; Prieto-Torres et al., 2018). Orogenic processes significantly affect landscape structure, promoting various speciation events due to their heterogeneous effect on climate, topography, and resource distribution, such as food availability (Gallen, 2018; Graham et al., 2014; Hoorn et al., 2010). For instance, isolated areas with high topographic complexity are often associated with an increase in speciation rates (Igea & Tanentzap, 2021; Steinbauer et al., 2016), as they form physical barriers that restrict connectivity between populations, limiting gene flow between them and favoring divergence and population structure through processes such as allopatric speciation (Gutiérrez-Pinto et al., 2012). Likewise, climatic gradients linked to elevation promote greater niche availability and heterogeneity, facilitating diversification driven by natural selection (Badgley et al., 2017; Love et al., 2023).

High mountain habitats are areas of special interest in the study of biogeographical and evolutionary patterns, as they form discontinuous geographical systems that restrict dispersal and connectivity between populations or species, similar to oceanic islands (Cerca et al., 2023; Lomolino et al., 1989; Sarthou et al., 2001). These secluded mountain peaks or mountain ranges isolated by inhospitable lowlands have been called “sky islands” (Nistelberger et al., 2015b; Ober & Connolly, 2015; Taylor, 1940).

The study of these systems has allowed the identification of common phylogeographic patterns in diverse taxa (Cox et al., 2014; Grismer et al., 2015; Love et al., 2023; Nistelberger et al., 2015a; Ober & Connolly, 2015; Robin et al., 2010). Some species of Afroalpine flora and the mountain reptile *Sceloporus jarrovii*, which are distributed in the sky islands of eastern Africa and southeastern Arizona, respectively, have experienced drastic population declines and, in some cases, local extinctions (Brochmann et al., 2022; Wiens et al., 2019). Such demographic events are attributed to the limited ability of these species to disperse and adapt in response to significant climatic fluctuations, such as glaciations, which caused the loss or reduction of high mountain habitats (Brochmann et al., 2022; Wiens et al., 2019). Therefore, demographic changes, characterized by repeated population bottlenecks and local extinctions, also appear to be a common pattern in sky islands (Love et al., 2023). Few studies have been conducted on sky islands in the Neotropical region, specifically in Colombia; most of them focused on plant species endemic to the páramo (Flantua et al., 2019; Sklenář et al., 2014; Valencia et al., 2020; Vásquez et al., 2016). These ecosystems are located between 3,000 and 5,000 meters above sea level (m.a.s.l.) across the Andean mountain range (Zapata et al., 2021). Research in this area has evaluated variation in connectivity of these high mountain ecosystems during substantial climatic changes, such as glacial periods, as well as how geographic isolation intervals in general led to divergence (Flantua et al., 2019).

Because sky islands create highly isolated, fragmented environments across sharp altitudinal gradients, they expose populations to abrupt ecological shifts that can fundamentally accelerate evolution and promote rapid speciation. Unlike standard allopatric divergence, rapid speciation occurs over remarkably compressed ecological timescales. This process can be heavily influenced by a number of factors, including ecological opportunity (such as ecologically based selection, local adaptation and migration) (Hendry et al., 2007; Momigliano et al., 2017), population dynamics (like behavioral traits and population size) (Veron et al., 2019) and intrinsic genomic architecture (like inversions, recombination, epistasis) (Campbell et al., 2018). For instance, strong divergent selection on specialized microhabitat preferences or assortative mating can drive partial reproductive isolation within merely dozens to a few hundred generations (Hendry et al., 2007; Nosil et al., 2009). Alternatively, internal genomic mechanisms can drastically accelerate this process; random macro-mutations, such as chromosomal rearrangements or inversions, can instantly establish physical barriers to recombination across the genome (Campbell et al., 2018), while a high mutation rate or positive selection on key reproductive loci can predispose certain lineages toward rapid speciation (Harvey et al., 2019; Oliver et al., 2009).

Within the northern Andes, the Sierra Nevada de Santa Marta (SNSM) represents a particularly compelling and unique mountain massif to investigate this dynamics due to the high endemism of species residing there, as well as the climate variability. The SNSM is not part of the Andean mountain range complex or a discontinuous set of sky islands. Rather, it constitutes an isolated geographical unit independent of the other mountain ranges of the country. Furthermore, it is located in northern Colombia, at the intersection of important tectonic faults that give it a complex geological history (Montes et al., 2010; SGC, 2019). It is the highest coastal mountain in the world, with an elevation of nearly 5,800 meters above sea level (Idárraga-García & Romero, 2010; Wiedemann, 2016) and is geographically isolated from the main mountain range system, having separated from the central mountain range approximately 30-40 million years ago (Piraquive et al., 2022; Tschanz et al., 1974). In terms of biodiversity, the SNSM is recognized for its highly heterogeneous ecosystems and vegetation cover ranging from savannas, swamps, and tropical rainforests, among others (Cavelier et al., 1998; Duran-Izquierdo & Olivero-Verbel, 2021). It also exhibits a high rate of endemism across several taxonomic groups, notably in amphibians (Granda-Rodríguez et al., 2014; Lorenzo et al., 2016), as shown by the presence of four morphospecies of the genus *Atelopus*, commonly referred to as harlequin frogs, which correspond to independent mitochondrial lineages (Lötters et al., 2025; Plewnia, Vega-Yánez, et al., 2026). Nonetheless, this taxonomic status has not yet been verified by other genomic regions.

The *Atelopus* genus is of great biological interest, mainly due to its high diversity in the Neotropics, with approximately 70 to 100 described amphibian species (Rueda Solano et al., 2016). They are generally distributed at altitudes above 1,500 meters above sea level, mainly found in northern South America and southern Central America (Lötters et al., 2011; Noonan & Gaucher, 2005). They are known for their aposematic coloration, which alerts predators of their toxicity, a trait shared by numerous species (Pearson & Tarvin, 2022). Nonetheless, research indicates a drastic decrease in population numbers for most species in the genus, with multiple extinction events of at least 30-40 species (Lampo et al., 2017; Lötters et al., 2023; Tapia et al., 2017). Despite this, studies have primarily focused on the genus’s ecological habits, behavior, and population monitoring in relation to the main factors driving its demographic decline (McCaffery et al., 2015).

The most complete phylogeny was inferred using both mitogenomics and short mitochondrial markers, which included the broadest sampling of species and geographic distribution yet seen of this genus (Lötters et al., 2025; Plewnia, Vega-Yánez, et al., 2026). This analysis shows that the SNSM species are a sister clade to all other *Atelopus* species, which are grouped into three main clades: the Andean-Venezuelan, the Andean-Chocó-Central American, and the Amazonian. Pattern that was also recovered in previous phylogenies (Lötters et al., 2011; Ramírez et al., 2020). Although significant progress has been made in the study of this genus, this research is not focused on understanding the speciation process of these toads, their historical demography and conservation status, specifically those in the SNSM region, which is included as one of the areas where the most recent common ancestor of all *Atelopus* spanned (Plewnia, Vega-Yánez, et al., 2026).

In particular, a correspondence among mitochondrial lineages does not necessarily imply that the same signal persists in nuclear DNA. The nuclear genome must be examined to definitively establish whether the lineages identified in the maternal line are maintained across other genomic regions, and to determine if speciation occurred (or is occurring) in the presence of gene flow (Long et al., 2026). Furthermore, the effect of barrier loci against genetic exchange (Stankowski & Ravinet, 2021) may be more consequential than, or act in along with, orogeny during the speciation process (Westram et al., 2022). This remains a crucial factor to be determined in the *Atelopus* of the SNMS.

Despite the interest for this genus there is notable lack of research on its phylogeography, behavior, ecology, speciation and demographic history specifically in the populations endemic to the Sierra Nevada de Santa Marta (SNMS). This is mostly due to difficult terrain access because of the presence of armed conflict in the region, as well as steep slopes and dense vegetation that hinders access to sampling areas. The SNSM comprises four endemic *Atelopus* species; *A.laetissimus*, *A.nahumae*, *A.carrikeri* (census Lotters et al. (2025)) the population sampled in this study could be *A.leoperezii*) and *A.arsyecue*. Although these species exhibit differences in morphology, few is known about their behaviour, distribution and ecology. For example, *A. carrikeri* described by Ruthven (1916), has five distinct color morphs (Rueda-Solano, 2011), is distributed above 3,000 m a.s.l. and prefers open, rocky areas with sparse vegetation within the páramo ecosystem (Rueda-Solano, 2011). On the other hand, regarding *A. laetissimus*—the most studied morphospecies—exhibits “mate-guarding” behavior, which involves prolonged amplexus (∼1 month) as an intense strategy of intrasexual selection (Rueda-Solano et al., 2022). The diversity of their acoustic signals has also been documented in both males and females in relation to the courtship process (Rueda-Solano et al., 2020). Furthermore, its reproductive effort in terms of energy spent has been quantified, alongside behavioral studies on its predation and habitat preferences (Rocha Usuga et al., 2017). In contrast, the morphospecies *A. arsyecue* remains poorly studied. Finally, each morphospecies tadpoles exhibited conserved coloration patterns and recognizable morphological differences, such as total length (J. Pérez-Gonzalez, 2017). Sexual dimorphism is present in *A. laetissimus*, *A. nahumae*, and *A. carrikeri*, where males are generally smaller (Sánchez-Ferreira, 2019). Only two cases of interspecific amplexus have been documented: one between the allopatric morphospecies *A. laetissimus* and *A. carrikeri* (J. L. Pérez-Gonzalez et al., 2017) and another between the sympatric *A. laetissimus* and *A. nahumae* (J. L. Pérez-Gonzalez et al., 2020). However, whether these encounters produced viable hybrids is unknown.

The lack of information in different aspects of this toads’ biology is concerning because factors such as habitat loss or modification (Gómez–Hoyos et al., 2020; McCaffery et al., 2015), changes in average temperatures and ecosystem characteristics (Kiesecker et al., 2001), and infection by the pathogenic fungus *Batrachochytrium dendrobatidis* - *Bd* (Flechas, Sarmiento, & Amézquita, 2012; Flechas, Sarmiento, Cárdenas, et al., 2012; Gass & Voyles, 2022; Scheele et al., 2019), known to cause a highly lethal chytridiomycosis, have decimated *Atelopus* populations whose genetic diversity remains unknown (but see Plewnia et al. (2026) as an exception in the SNMS).

Consequently, it is crucial to strengthen the study of geographically complex regions such as the SNSM, as a sky island that harbors a significant quantity of endemic and endangered species, including certain species of the genus *Atelopus*. This research aims to confirm the taxonomic status of *Atelopus* species found in the SNSM, determine if their differentiation occurs in the presence of gene flow, and examine their genetic diversity and structure, as well as their evolutionary and demographic history. Finally, the estimation of effective population sizes (N_e_) over time, in contrast to available censuses, will be useful to evaluate the current state of genetic diversity within each group and, ultimately, its susceptibility to decline and risk of extinction.

## Material and methods

### Taxon sampling

The “Fundación Atelopus” (https://www.fundacionatelopus.org/) collected samples across the Magdalena, Cesar, and La Guajira departments in five locations within the SNSM (Figure 1). The specimens were collected at elevations ranging from approximately 1,500 to 3,600 m a.s.l., which is the reported preferred distribution range for the four morphospecies. Additional specimens were collected on the Colombian Pacific coast, belonging to *A. spurrelli*, and *A. fronterizo*, located near the border with Panamá and the Caribbean coast. These later species were used as the outgroup.

**Figure 1.**
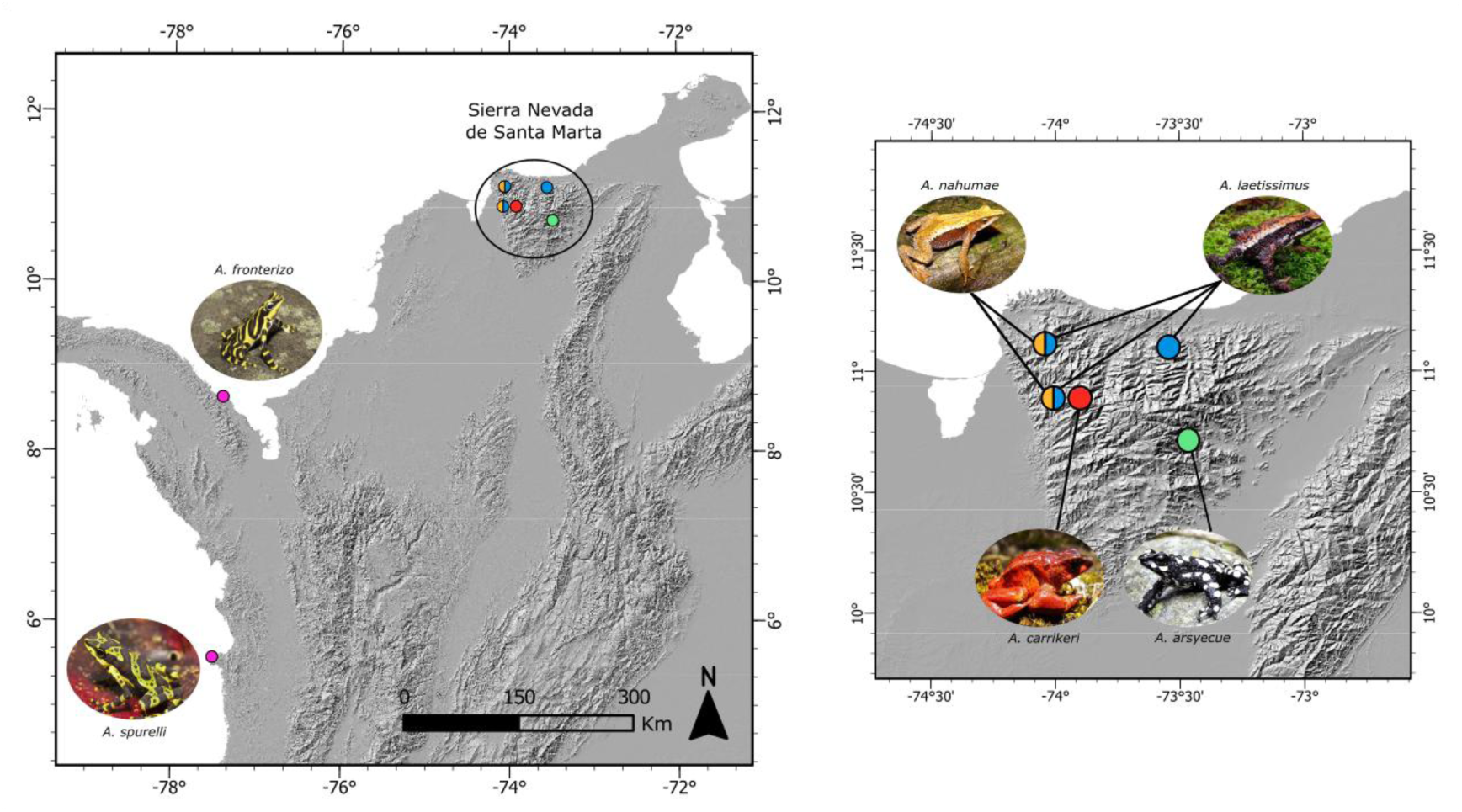
Sampling locations. The species selected as the outgroup (*A. spurrelli* and *A. fronterizo*) were found along the western coast of Colombia. The colors represent the different sampling locations for each of the respective morphospecies: Blue: *A. laetissimus*, green: *A. arsyecue*, orange: *A. nahumae*, and red: *A. carrikeri*. *A. laetissimus* and *A. nahumae* share two sampling locations (represented by the two-colored circles). Photos courtesy of Fundación Atelopus.

Tissue collection was made by making a transversal incision in the outer phalanx of the right hind leg of each sample. A surgical sealant was applied to prevent potential infections. 95 individuals were collected in total, in detail: 89 individuals belonging to the four morphospecies present in the SNSM and 6 individuals from the two continental species (Table S1; Figure 1). The tissues were preserved in 2-ml cryovials with 90% alcohol and stored at -80°C. The sex of each individual was recorded with the assistance of experts from Universidad del Magdalena, and the morphospecies were identified based on morphological characteristics (Rueda Almonacid et al., 2004).

### DNA Extraction and Next-Generation Sequencing (NextRAD)

DNA was extracted using the SeraPure magnetic bead method (Rohland & Reich, 2012). The tissue was digested using a lysis buffer (180 μl), 20 μl of proteinase K was added (by inverting the tube) before incubation at 56 °C for 24 hours. The DNA concentration and purity were quantified by fluorometry (Qubit, Thermo Fisher) and Thermo Scientific™ NanoDrop, respectively. Additionally, the quality and integrity of the genomic DNA were verified in 1.5% agarose gels.

Genotyping libraries were constructed, and variable sites were identified through transposase-mediated DNA sequencing using the nextRAD method (Russello et al., 2015). Genomic DNA was fragmented using the Nextera reagent (Illumina, Inc.), which also ligated adapter sequences to the ends of the DNA fragments. These fragments were amplified over 27 cycles through the binding of primers that match to the adapter sequences and extend 10 nucleotides into the genomic DNA with the selective sequence “GTGTAGAGCC”, ensuring that only fragments beginning with a sequence hybridized to the selective primer sequence were efficiently amplified. NextRAD libraries were sequenced on a Novaseq 6000 system at the University of Oregon to generate 122-bp single-end reads. (ENA accession number xxxx-xxxx).

### Coverage relative to the Atelopus reference genome and the presence of mtDNA reads

Quality control of the obtained reads was performed using the FASTQC v0.11.9 (https://www.bioinformatics. babraham.ac.uk/projects/fastqc/) and MultiQC v1.6 (Ewels et al., 2016). The detected Nextera adapters were subsequently removed using the Trimmomatic v0.38 program (Bolger et al., 2014). To verify the presence of mitochondrial DNA reads, these were mapped to the mitochondrial genome of *Rhinella marina*, GenBank accession No. NC_066225.1, which is the most similar to the SNSM morphospecies, as referenced in the publication of the *Atelopus laetissimus* genome (Araújo et al., 2025). The reference mitochondrial genome was previously indexed using the BWA v0.7.17 software (Li, 2013). Subsequently, the reads were aligned and processed using Samtools v1.8 (Li et al., 2009), and the mapping results were summarized using the flagstat function.

In addition, the reads were mapped to the *A. laetissimus* genome (Araújo et al., 2025), NCBI accession No. JBGDNC00000000, to estimate the percentage of the reference genome covered by each individual. The assembled genome, consisting of 1,458 contigs corresponding exclusively to nuclear DNA, was downloaded and consolidated into a single FASTA file. Subsequently, the reference genome was indexed, and the reads were aligned using BWA v0.7.17 (Li, 2013) and Samtools v1.8 (Li et al., 2009).

We used the depth-a function in Samtools v1.8 (Li et al., 2009) to calculate the coverage percentage per individual, which reports the alignment depth at all positions in the reference genome. The coverage percentage was estimated as the proportion of positions in the reference genome with an alignment depth greater than zero, considering all evaluated positions.

### Filtering and SNP calling

For genotyping, custom scripts provided by SNPsaurus were implemented, following the method described in (Salgado-Roa et al., 2024). Using bbduk from the BBMap package (Bushnell, 2014), the adapters used in sequencing were removed, the ends of low-quality reads (<20) were trimmed, and a de novo reference assembly was generated with a total of 10 million reads, which were uniformly sampled across all individuals. Additionally, all reads with a depth greater than 600 and less than 6 were removed. The remaining reads were aligned against each other to identify allelic loci and collapse them into a single representative haplotype. Finally, the reads were mapped to the obtained assembly using an alignment identity threshold of 0.90 via bbmap (BBMap package). SNPs were called using BBMap’s callvariants with a depth of 15x (Bushnell, 2014).

Finally, the resulting VCF file containing the variant calls (SNPs) was used as a basis and filtered using VCFtools v.0.1.16 (Danecek et al., 2011) following (Salgado-Roa et al., 2024). Only those loci present in at least 70% of the individuals, as well as biallelic SNPs, were retained, for a total of 95 individuals and 32,008 SNPs kept.

### Phylogenetic inference and species delimitation

Following the filtering recommendations of (Hemstrom et al., 2024), reads with > 60% missing data were removed using VCFtools v.0.1.16 (Danecek et al., 2011) as well as SNPs that did not meet the Hardy-Weinberg equilibrium assumption. SNPs that exhibited linkage disequilibrium were removed using a custom script ldPruning.sh (https://github.com/joanam/scripts). A total of 83 individuals remained, 79 corresponding to the morphospecies and 4 to the outgroup, along with 899 SNPs.

The phylogenetic relationships between the morphospecies and outgroups were determined using maximum likelihood analysis with RAxML v.8.2.10 (Stamatakis, 2014), under the GTRGAMMA substitution model, which allows for heterogeneity in the substitution rate across sites. Node support was obtained using 1,000 bootstrap replicates, applying Lewis’s verification bias correction (Lewis, 2001), a function implemented in RAxML (--asc-corr=lewis).

The IQ-TREE 2 software (Minh, Schmidt, et al., 2020) was also implemented using the ModelFinder function (Kalyaanamoorthy et al., 2017) to determine the best phylogenetic model that fit the data; in addition, validation bias correction (MFP+ASC) (Lewis, 2001) was included. Branch and node support (-alrt and abayes in IQ-TREE) were used, with 1,000 replicates in each case. The SH-aLRT method was implemented for branch support (Anisimova & Gascuel, 2006; Guindon et al., 2010). Also, a Bayesian version of aLRT (aBayes) was used (Anisimova et al., 2011). For node support, the ultrafast bootstrap (UFBoot) method was implemented (Hoang et al., 2018).

In addition, SVDQuartets (Chifman & Kubatko, 2014) was used, a multi-species coalescent phylogenetic inference approach implemented in PAUP* v4.0a169 (Cummings, 2004); individuals were first assigned to a suggested species, and the species tree was constructed using the quartet method (Chou et al., 2015); the nodes were supported by 10,000 bootstrap replicates. Additionally, a phylogenetic network was estimated in SplitsTree using the Neighbor-Net algorithm (Huson & Bryant, 2006).

Finally, we implemented the Bayesian factor-based species delimitation method (BFD*) (Leaché et al., 2014) using the BEAST v.2.7.7 software (Bouckaert et al., 2019; Bryant et al., 2012), coupled with SNAPPER (Stoltz et al., 2021), which uses multispecies coalescence, allowing for the estimation of species trees based on SNPs. The prior parameter values were set according to the recommendations in A. Leaché’s SNAPPER implementation guide (https://taming-the-beast.org/tutorials/BFD_snapper_tutorial/BFD_snapper_tutorial.pdf). The coalescence rate distribution was assigned based on the estimated height of the species tree, which was calculated using the average of the maximum divergence between groups estimated using MEGA v12 (Kumar et al., 2024) divided by two. This was modeled using a gamma distribution with a mean of 20 (α = 2, β = 10). The other values were kept at their default settings. We developed three different models for species delimitation; marginal probability values were estimated for each model using path sampling with 48 steps, a chain length of 100,000 generations, and a pre-burn-in of 10,000. Bayes factor (BF) scores were calculated between each alternative model using their marginal likelihood values (Leaché et al., 2014). Convergence of the chains and verification of ESS values greater than 200 were assessed in Tracer v1.7.2 (Rambaut et al., 2018). The consensus tree was obtained using TreeAnnotator v.2.7.7 (Bouckaert et al., 2014), visualized with FigTree v.1.4.4, and statistics on the distribution of topologies were obtained using TreeSetAnalyser, part of the BEAST v.2.7.7 software (Bouckaert et al., 2019).

Since the inference methods used resulted in different topologies and there is no consensus regarding the phylogenetic relationships among the morphospecies, we performed a topology test in IQ-TREE 2 (Minh, Hahn, et al., 2020). This comparison was supplemented primarily by the Approximately Unbiased (AU) test (Shimodaira, 2002) and other secondary tests (Kishino et al., 1990; Kishino & Hasegawa, 1989; Shimodaira & Hasegawa, 1999; Strimmer & Rambaut, 2002) to statistically evaluate the level of support for each topology in relation to the data.

### Summary statistics of intra- and inter-morphospecies diversity

Nucleotide diversity (π) and Tajima’s D per tag per morphospecies were calculated using the *Genopop* package in R (Gurke, 2023). Heterozygosity and inbreeding coefficient per site per individual were estimated using VCFtools v.0.1.16 (Danecek et al., 2011). These calculations were performed using the initial VCF file containing the 32,008 SNPs.

To establish and evaluate patterns of genetic differentiation among morphospecies, we applied a more conservative filtering by removing reads with >50% missing data. Subsequently, SNPs that did not meet the Hardy–Weinberg equilibrium assumption and those in linkage disequilibrium were removed. For these procedures, VCFtools v.0.1.16 (Danecek et al., 2011) and the ldPruning.sh script (https://github.com/joanam/scripts) were used, respectively, resulting in a total of 78 individuals and 1,024 SNPs (one SNP per tag).

### Analysis of genetic clustering and ancestry

Initially, a Principal Component Analysis (PCA) was performed using the PLINK package v.1.90 (Purcell et al., 2007). Subsequently, we used the *adegenet* package (Jombart & Ahmed, 2011) for several downstream analyses: The number of genetic clusters (K) was estimated using the *find.clusters* function, retaining 60 components, and the a Discriminant Analysis of Principal Components (DAPC) was performed selecting the optimal number of retained components via cross-validation using the *xvalDapc* function. Finally, a MANOVA analysis was performed using the *manova* function, employing the DAPC “linear discriminants” as dependent variables and the morphospecies as the independent variable.

FastSTRUCTURE and ADMIXTURE v.1.3.0 were used to determine the number of genetic clusters (K) that best explains the genetic variation within a range of 1 to 10. This was done using the *chooseK.py* function in fastSTRUCTURE (Raj et al., 2014) and cross-validation in ADMIXTURE (Alexander et al., 2009). In addition, the population structure and the haplotype-based co-ancestry matrix were inferred using the RADpainter package and the posterior probability of clustering was estimated using fineRADstructure (Malinsky et al., 2018). For this, clustering was done with 100,000 MCMC iterations (sampling every 1,000) and a 100,000-iteration burn-in. The final fineRADstructure tree was constructed using 10,000 iterations.

Finally, the F_st_ and D_xy_ statistics were calculated by tag for pairs of morphospecies, using the R package *Genopop* (Gurke, 2023) and the VCF file containing 32,008 SNPs. The D_a_ statistic was calculated using the formula defined by Nei & Li (1979).

### Historical demography and genetic flow patterns

Demographic history was reconstructed using the same VCF employed for the structure analyses. We used the Genetic Algorithm for Demographic Model Analysis-GADMA2, which utilizes the joint site allele frequency spectrum (JSFS) generated with easySFS (Gutenkunst et al., 2009). This software infers the best demographic model based on the maximum likelihood value obtained from the simulation of the joint site frequency spectrum that most closely approximates the observed JSFS. Additionally, GADMA2 also considers the calculation of the Akaike Information Criterion (AIC) to avoid model overfitting through excessive optimization of the parameters used in the simulations (Noskova et al., 2020, 2022). These models provide estimates for divergence times, historical changes in effective population size, and potential migration events.

Because GADMA2 does not require a predefined model structure, the number of generated models varies. We performed four distinct simulations, each with three populations, to evaluate all possible groupings among the four morphospecies; this approach was taken as GADMA2 is optimized for handling up to three populations (Noskova et al., 2020, 2022). Each simulation was replicated three times with a [1, 1, 1] initial structure. The mutation rate was set to 1.2 × 10⁻⁹ substitutions per site per year, based on the estimate made for the amphibian genus *Eleutherodactylus* (Crawford, 2003), and a generation time of 2 years (Jorge et al., 2022). Additionally, we selected the Powell method for optimizing the local search algorithm and the *moments* engine (Jouganous et al., 2017), as these have been reported to be the most effective (Noskova et al., 2020).

We ran two scenarios for each of the four simulations. The first scenario used a model of isolation and divergence of morphospecies in the complete absence of migration; the second allowed for asymmetric migration events throughout the entire simulated time period. All other parameters were kept at their default values in both scenarios. Each simulation was run with three replicates. Although the isolation scenario minimized the AIC values, the site frequency spectrum approximated by these models differed considerably from the observed JSFS, which was reflected in a greater dispersion of the Anscombe residuals. Because of this, the models that allowed migration between morphospecies were chosen.

Based on the migration scenarios identified by GADMA2, we tested for potential introgression among the four morphospecies using the *D-* and *f4*-statistics implemented in Dsuite v0.5 (Malinsky et al., 2021), based on the ABBA-BABA patterns described by Patterson et al. (2006). The significance of the *D-* and *f4* statistics was assessed using their respective Z-scores, calculated via a standard block-jackknife procedure; furthermore, significance of the tests was verified using the p-value (Malinsky et al., 2021). To assign the ancestral allele, the species *A. fronterizo* and *A. spurrelli* were used as outgroups for all possible combinations of three species. Subsequently, we removed the outgroup using the *Dquartets* function to evaluate the four morphospecies present in the SNSM as an unrooted tree with an arbitrary assignment of the ancestral allele. Both analyses were compared with each other and with the previously obtained ancestral groupings.

Additionally, to determine whether the presence of shared polymorphisms among the morphospecies could be explained by incomplete lineage sorting (ILS), we estimated the probability of this phenomenon occurring following Knief et al. (2024). This estimation is based on the principles of coalescence theory; therefore, the probability is derived as a function of the divergence time between species (τ = (D_a_/π) (Blumer et al., 2025; Wiuf et al., 2004), their current effective sizes (N_1_ and N_2_), and the effective size of the ancestral population (N_a_) (Knief et al., 2024), parameters that were previously inferred using GADMA2 and the *Genopop* package in R (Gurke, 2023). We calculated the probability using the following expression, which approximates the lower bound of the fraction of shared polymorphisms relative to the total number of polymorphisms present in the sampled lineages; this assumes neutrality and the absence of migration between species (Knief et al., 2024).

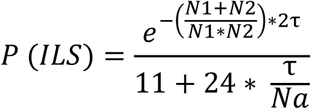

### Verification of the change in effective size

We used Stairway Plot v.2 (Liu & Fu, 2020) to model historical fluctuations in the effective population size of each morphospecies. This analysis aimed to validate the demographic trends observed in the GADMA2 models. The input SFS was generated via easySFS, and all calculations employed the same mutation rate (1.2×10⁻⁹ per site per year) and generation time of (2 years) used in the prior demographic analyses (Crawford, 2003; Jorge et al., 2022; Lau et al., 2020).

Finally, since *A. laetissimus* is the only morphospecies for which population censuses have been conducted in recent years, the proportion of the population contributing to reproduction was calculated. The *N_e_/N_c_* formula was used, where *N_e_* corresponds to the effective population size calculated by Stairwayplot and *N_C_* to the population census (Clarke et al., 2024). For the population census, data were recorded from two populations of *A. laetissimus*: the locality known as San Lorenzo, with 342 individuals (308 males and 34 females), and San Pedro, with 282 individuals (244 males and 38 females), for a total of 632 individuals (Atelopus Foundation, unpublished data).

## Results

### Coverage relative to the Atelopus reference genome and presence of mitochondrial DNA reads

On average, NextRad reads covered 2.22% ± 1.10% of the *A. laetissimus* genome per individual. Depending on the specific filters applied (see Material and methods), coverage ranges between 0.0031% and 0.0035% when expressed as the number of SNPs present in the data. Because the *A. laetissimus* genome assembly does not contain mitochondrial scaffolds, we used *Rhinella marina* as a reference to verify the presence of mitochondrial reads in the data. This revealed low mtDNA content in two individuals: one *A. carrikeri* (0.005%) and one *A. fronterizo* (0.02%).

Based on these results, the *A. fronterizo* individual was excluded from the study due to its mitochondrial DNA percentage and the availability of four other individuals to serve as an outgroup. In the case of *A. carrikeri*, we performed preliminary analyses both including and excluding the individual with mtDNA traces. Since its exclusion did not produce changes in the results, this individual was retained in the final analysis.

### Phylogenetic inference and species delimitation

Both maximum likelihood phylogenetic trees and species delimitation methods recover each of the morphospecies as monophyletic with high support (> 90%; Figure 2; Figures S1–S3), additionally, *Atelopus* morphospecies native to the Sierra Nevada de Santa Marta (SNSM) were recovered as a single monophyletic group, clearly divergent from the outgroup taxa (100%; Figure 2; Figures S1 and S3). The 4-species model had the highest marginal likelihood and received decisive support over the others model based on the BF values. However, phylogenetic relationships among the morphospecies were discordant across methods. While *A. laetissimus* and *A. arsyecue* appear as sister species with low bootstrap support in RAxML and SVDQuartest (< 54%; Figures 1S and S3) this grouping is supported in IQ-TREE 2 (55% in SH-aLRT, 88% in UFBoot and 100% in aBayes; Figure 2). On the other hand, *A. carrikeri* was recovered as a sister taxon to the aforementioned clade in IQ-TREE 2 (70% in SH-aLRT, 64% in UFBoot, and 99% in aBayes; Figure 2), whereas *A. nahumae* was placed as a sister taxon to the clade formed by *A. laetissimus* and *A. arsyecue* in the trees generated by RAxML and SVDQuartest (94% bootstrap in RAxML and 39% in SVDQuartest; Figures S1 and S3).

**Figure 2.**
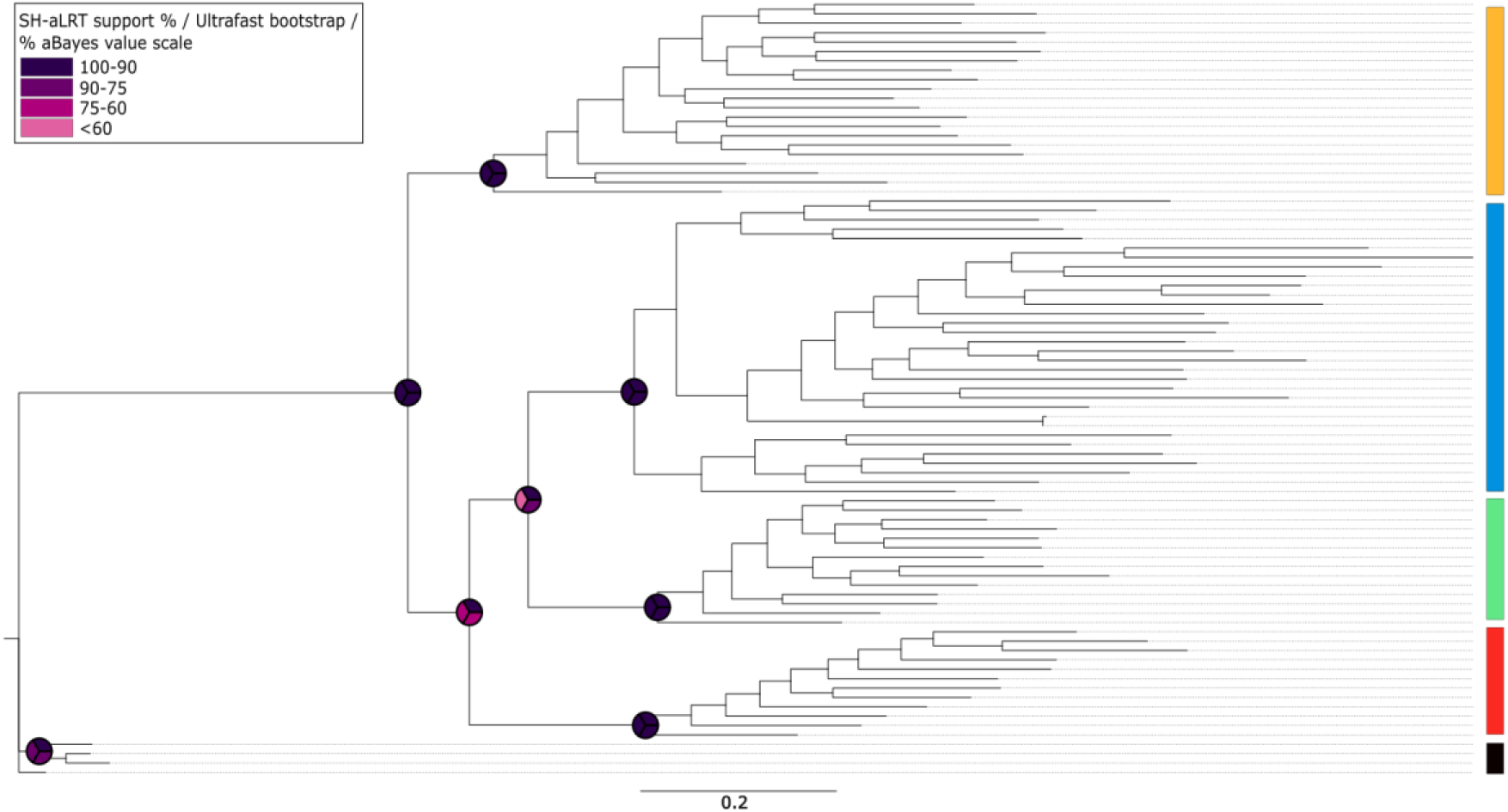
Maximum likelihood phylogenetic tree inferred with IQTREE 2. Colored circles at the nodes indicate node support values for SH-aLRT, UFBoot, and aBayes, ranging from purple (100% support) to pink (< 60% support). Morphospecies are identified by color: blue: *A. laetissimus*, green: *A. arsyecue*, orange: *A. nahumae*, red: *A. carrikeri*, and black: Outgroup.

The phylogenetic tree inferred using SNAPPER groups *A. nahumae* and *A. carrikeri* as sister species (posterior probability of 0.99; Figure S2), with *A. laetissimus* as a sister species to this pair (posterior probability of 1), while *A. arsyecue* was placed as the sister species to the clade comprising the other morphospecies (posterior probability of 1; Figure S2).

The SplitsTree distance network clearly partitioned the four morphospecies into distinct groups. However, the presence of multiple reticulations among the clusters suggests potential hybridization events and/or ILS (Figure S4). These patterns are consistent with the topological discordance observed across our phylogenetic analyses. Specifically, only 72.08% of the 1,929,501 quartets evaluated in SVDQuartets are consistent with the final topology inferred by this method and described above (Figure S3).

The topology test confirmed that the topology with the highest support was the one inferred using maximum likelihood with IQ-TREE 2, which also yielded the best log-likelihood value (AU test, p > 0.05; Table 1 and Table S2). Furthermore, compared to the other phylogenetic inference methods, this topology generally exhibited the highest branch support values for the relationships among the morphospecies (Figure 2).

**Table 1.**
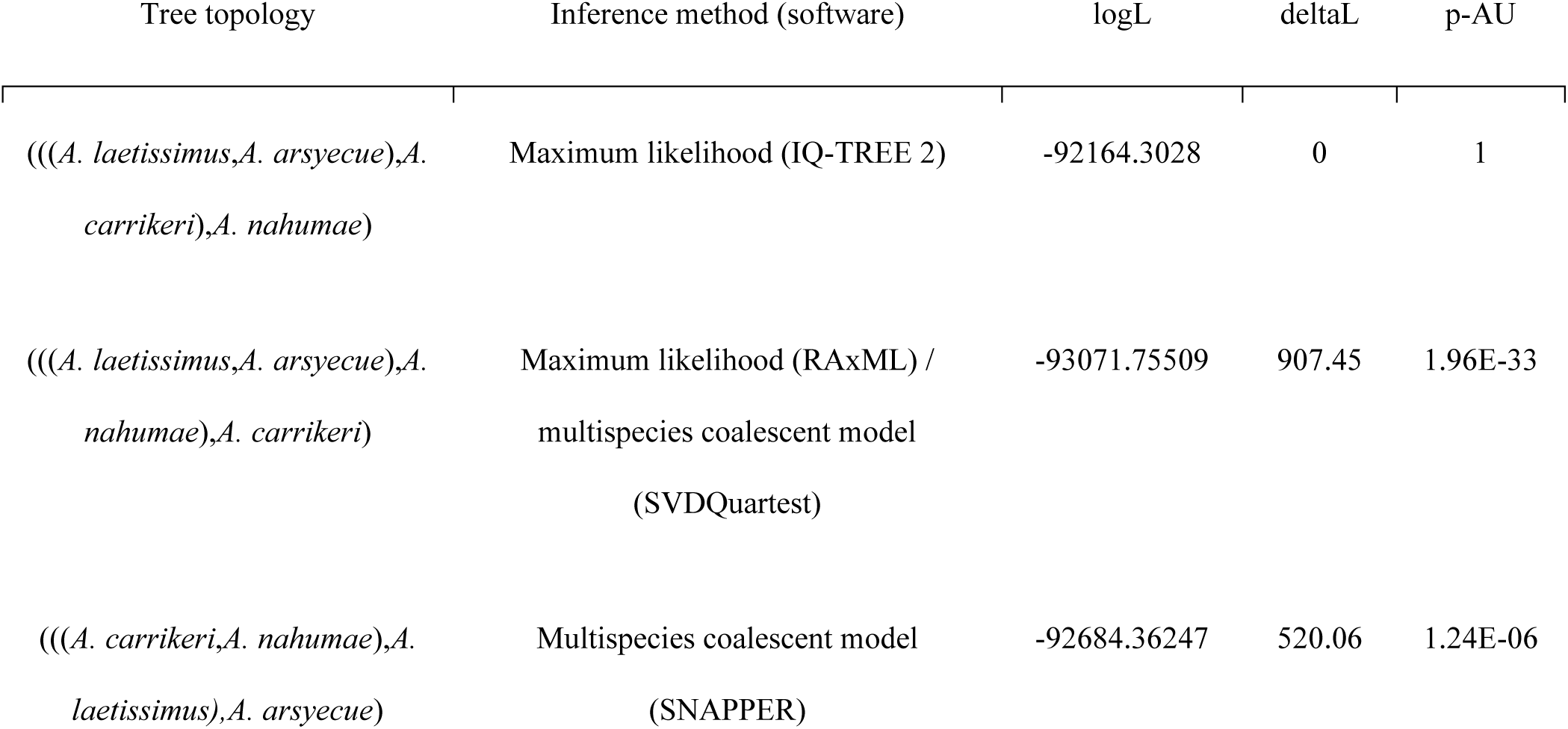
Log-likelihood values and Approximately Unbiased (AU) test results for the evaluated tree topologies. Results for all other topology tests are provided in Supplementary Table 1.

### Summary of intra- and inter-morphospecies diversity statistics

Average nucleotide diversity among the four species ranged from 3.99×10⁻³ to 5.64×10⁻³; with the lowest value observed in *A. arsyecue* and the highest in *A. laetissimus* (Figure S5). All pairwise comparisons showed significant differences (p < 0.005; Figure S5). Regarding heterozygosity per individual, *A. laetissimus* had the lowest values, followed by *A. nahumae* (Figure S6). While *A. carrikeri* exhibited the highest heterozygosity, it was not significantly different from *A. arsyecue* (Figure S6). Aside from this specific comparison, all other pairwise comparisons were significant (p < 0.005; Figure S6).

Across all three genetic differentiation statistics (F_st_, D_xy_, and D_a_) the comparison between *A. laetissimus* and *A. arsyecue* showed the lowest values among all morphospecies (Figures S7–S9). In contrast, comparisons including *A. nahumae* showed the highest average D_xy_ and D_a_ values (Figures S8 and S9) excluding F_st_, where genetic differentiation between *A. arsyecue* and *A. carrikeri* is similarly high (Figure S7). While differentiation (F_st_) was lower for pairs including *A. laetissimus*, absolute measure of divergence (D_xy_) did not follow this trend; instead, D_xy_ values involving *A. laetissimus* were similar to other comparisons, being the highest values those between *A. laetissimus*-*A. nahumae* and *A. arsyecue*-*A. nahumae* (Figures S7 and S8). Finally, net divergence (D_a_) mirrored the F_st_ results where pairs including *A. laetissimus* showed smaller distances than those without (Figure S9). The average D_a_ values across all morphospecies fell below the lower threshold of the “gray zone of speciation” (0.5%; Figure S9). However, a substantial number of loci fell within the gray zone range (0.5% < D_a_ < 2%), and several loci exceeded the upper threshold (2%).

### Analysis of genetic clustering and ancestry

Principal Component Analysis (PCA) revealed that genetic diversity is distributed across four distinct groups, each corresponding to a SNSM species; the first two components accounted for 59.1% of the total variation (Figure S10). This clustering was consistent across DAPC, ADMIXTURE, and fastSTRUCTURE analyses (Figures S11–S13), which also identified the optimal number of genetic clusters to be K = 4, one for each species. MANOVA results indicated significant differences between these morphospecies-defined groups (p < 0.05). While K=4 was optimal, both ADMIXTURE and fastSTRUCTURE also suggested K=5 as a highly likely clustering level, though ADMIXTURE yielded similar CV values for K=3 and K=5 (Figure S14). At K = 5, both analyses identified a strongly structured population within *A. laetissimus*, separating individuals collected at the localities known as Serranía de las Cebolletas (municipality of Ciénaga-Magdalena) and Río Ancho (municipality of Dibulla-La Guajira), excluding the third locality of San Lorenzo (municipality of Santa Marta-Magdalena) (Figures S12-S13). However, at K = 3, ADMIXTURE groups *A. arsyecue* and *A. carrikeri* species into the same genetic cluster, which differs from fastSTRUCTURE, which groups *A. arsyecue* with *A. nahume*. Additionally, ADMIXTURE estimates a higher proportion of common ancestry among individuals of *A. laetissimus* and other species.

These structure analyses are partially consistent with the co-ancestry matrix, which exhibits a distinct population structure among the three *A. laetissimus* populations (lower left corner in Figure 3). Specifically, *A. laetissimus* population collected in the Serranía de las Cebolletas shows greater coancestry with *A. carrikeri* and *A. nahumae*, whereas the Río Ancho population shows greater coancestry with *A. arsyecue* (eastern side of the SNSM). This “admixture” pattern (∼20%) was consistently recovered in both fastSTRUCTURE (Figure S12) and ADMIXTURE (Figure S13). Furthermore, individuals of *A. laetissimus* are grouped by population in phylogenies, although these nodes received low support (Figure 2 and Figure S1). In contrast, no evidence of genetic structure or geographic grouping was found within *A. nahumae* (Figures 2-3 and Figures S1, S12–S13) .

**Figure 3.**
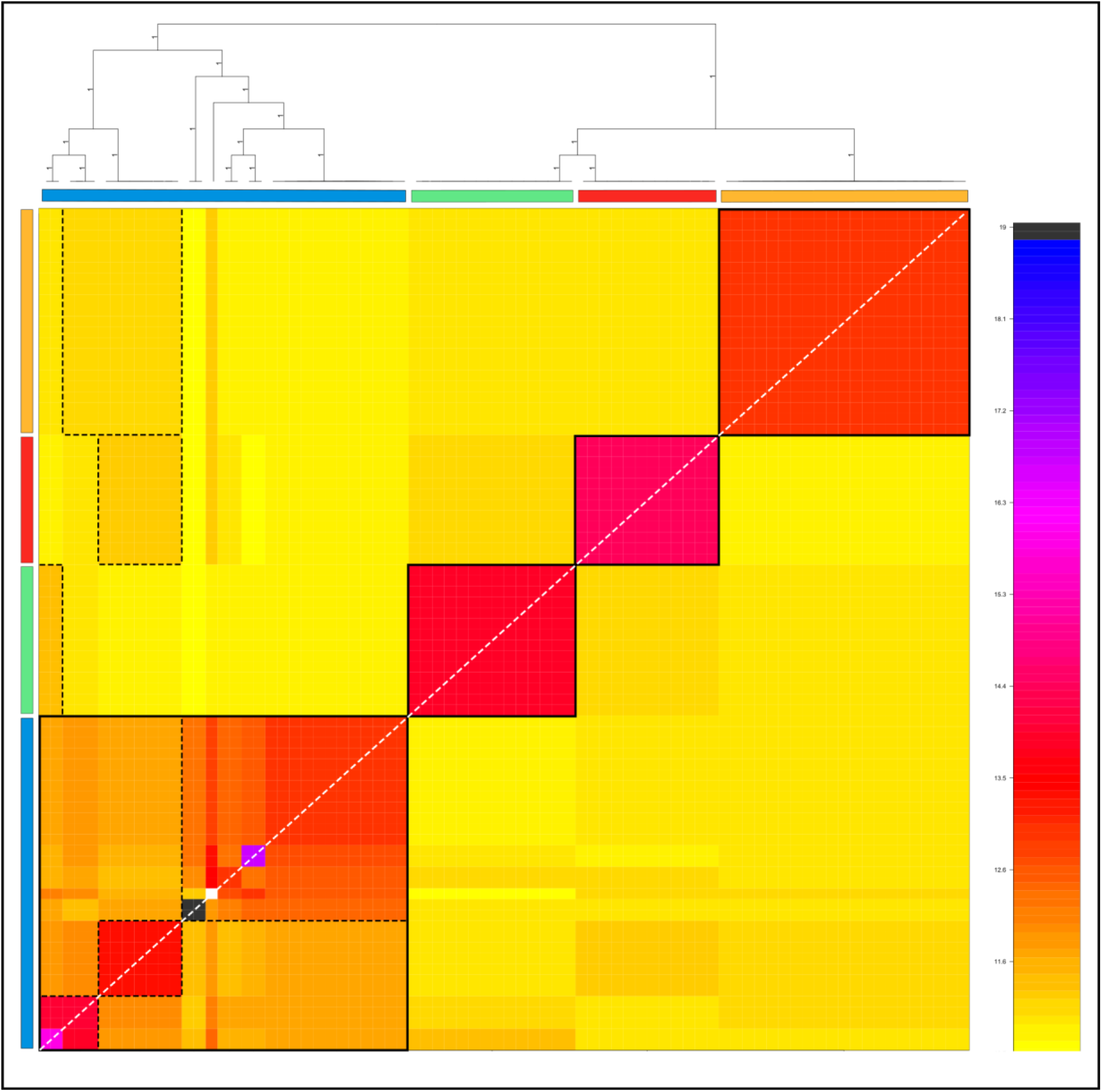
Co-ancestry matrix inferred by fineRADstructure. Solid black squares highlight the distinct population structure within each morphospecies, whereas dotted black squares highlight shared co-ancestry across populations and morphospecies. Species are identified by color: blue: *A. laetissimus*, green: *A. arsyecue*, orange: *A. nahumae* and red: *A. carrikeri*.

### Historical Demography and genetic flow patterns

The optimal demographic models inferred by GADMA2 for each of the three species combinations as inferred by the best topology tree showed consensus on several of the estimated demographic parameters (Table S3). Across all four models, the first divergence event from the ancestral populations (regardless of the species included in the analysis) was estimated to have occurred within the last 1.5 million years (1.410.917, 966.894, 977,695, and 849.695 years; Figure 4A–D). Subsequently, the second divergence event between lineages was recovered within less than 250,000 years (47.050, 48.554, 132.813, and 227.235 years; Figure 4B–D).

**Figure 4.**
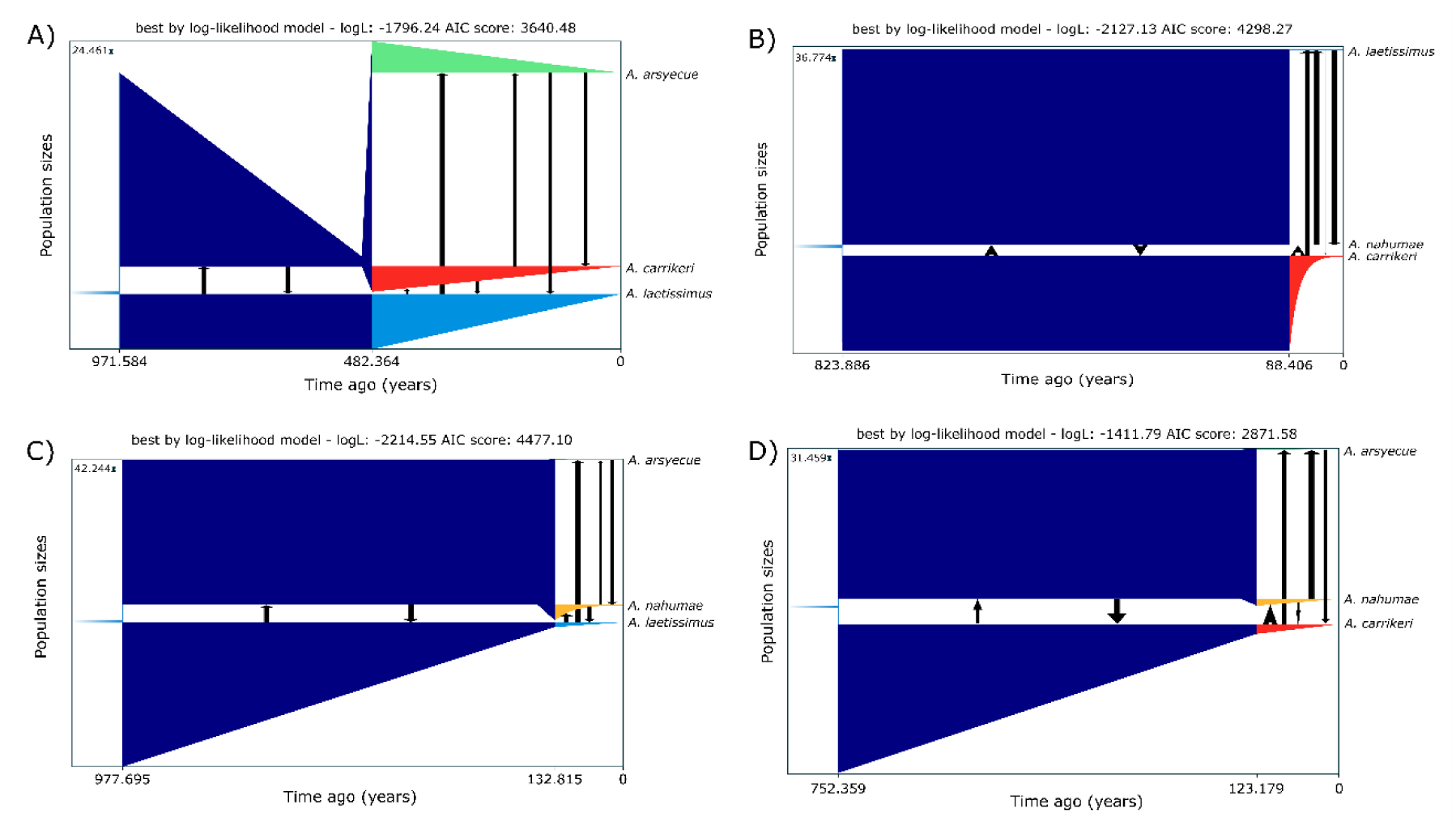
Best-fitting demographic models inferred by GADMA2 using the “moments” engine for four distinct three-species simulations: (A) *A. carrikeri*, *A. laetissimus*, and *A.arsyecue*. (B) *A. nahumae*, *A. carrikeri*, and *A. laetissimus*. (C) *A. nahumae*, *A. laetissimus*, and *A. arsyecue*. (D) *A. nahumae*, *A. carrikeri*, and *A. arsyecue*. Arrows indicate migration events and their direction, with arrow size proportional to the effective migration rate. Polygon size is proportional to the effective population size (N_e_) of ancestral populations and species. Species are identified by color: blue: *A. laetissimus*, green: *A. arsyecue*, orange: *A. nahumae* and red: *A. carrikeri*.

A drastic contraction in effective population size was observed across all species following the second divergence event, with reductions ranging from ∼92% to ∼99% (Figure 4A-D). In some cases, this decline began in the ancestral populations even prior to the split. Furthermore, the models recovered evidence of migration between both ancestral lineages and species (Figure 4A-D). These migration events were generally asymmetric, with high effective migration rates in most cases. Lastly, demographic modeling identified migration events among both ancestral populations and species (Figure 4A–D). These migration events were asymmetric between ancestral and contemporary populations, exhibiting high effective migration rates in most cases (Table S3).

In contrast to the high effective migration rates inferred by our demographic models, tests of shared site frequency (ABBA-BABA patterns) detected no significant signal of introgression across any of the evaluated species combinations (Tables S4 and S5), suggesting a scenario with no gene flow. Additionally, the estimated probability of ILS among the different species pairs was ∼9.08%, assuming neutrality and the absence of migration.

### Verification of the change in effective size

The Stairway Plot recovered a pattern consistent with GADMA2 simulations, showing a marked contraction in the effective population size of all four species over the last 200,000 years (Figure 5). Following this decline, *A. nahumae*, *A. arsyecue*, and *A. carrikeri* exhibited a demographic expansion starting at approximately 8,000, 40,000, and 75,000 years ago, respectively, with *A. carrikeri* showing the most pronounced recovery (Figure 5). Tajima’s D statistic provided additional evidence of possible bottlenecks, with a notable proportion of loci displaying values exceeding 2 (around 6.33% in *A. nahumae*, 9.24% in *A. laetissimus*, 5.36% in *A. carrikeri*, and 9.84% in *A. arsyecue*; Figure S16). Conversely, the inbreeding coefficient F_(HOM)_ showed no signs of recent bottlenecks, although individuals of *A. laetissimus* displayed lowest values for this statistic (mean = 0.21; p < 0.05) (Figure S17). Finally, the estimated *N_e_/N_c_* ratio for *A. laetissimus* was 0.54 was obtained, suggesting that roughly half of the population actively contributes to the breeding pool.

**Figure 5.**
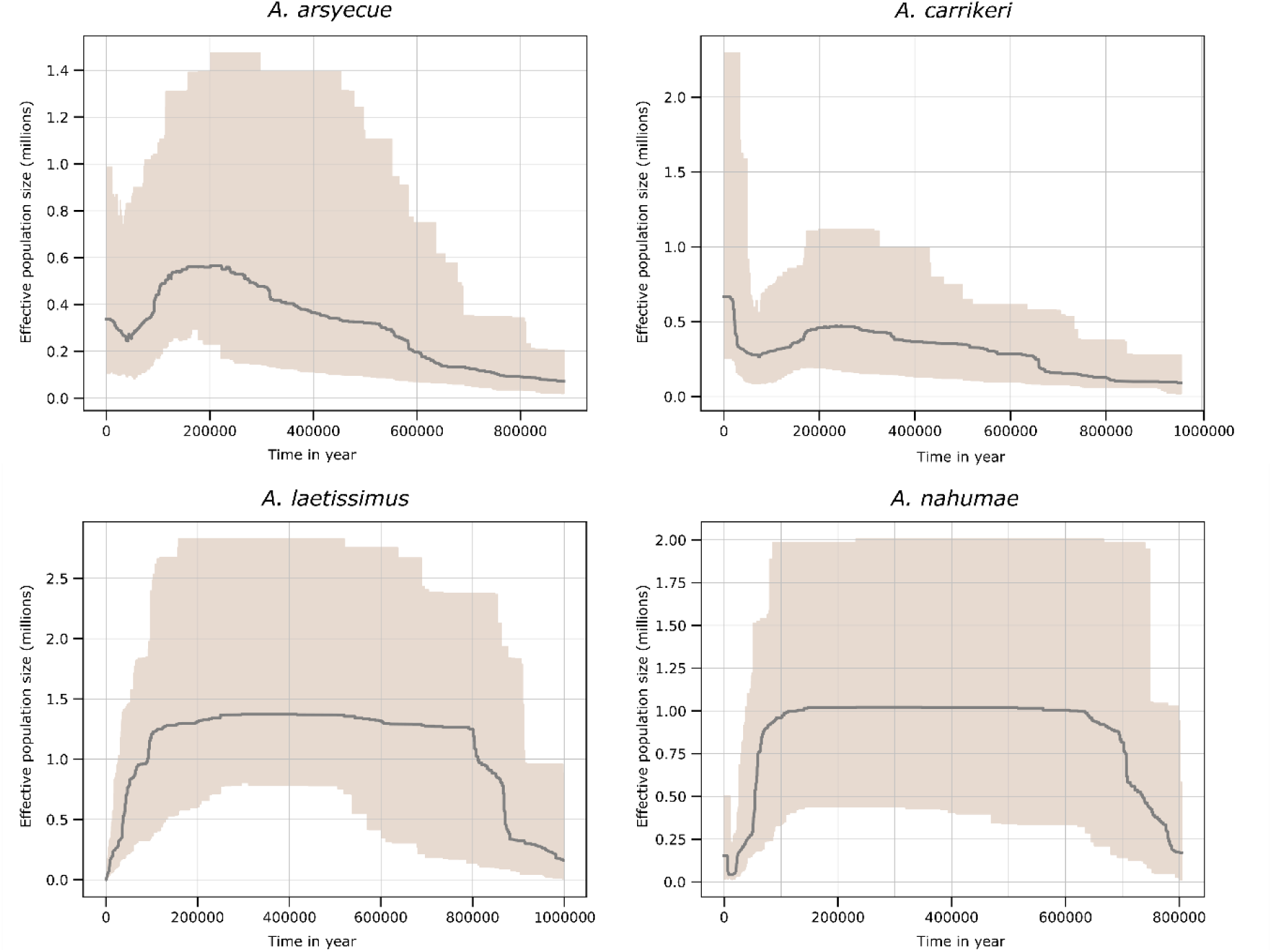
Stairwayplot estimates of the effective population size (N_e_) over time for the four species present in the SNSM. Calculated based on the site frequency spectrum (SFS) specific to each species. The shaded areas represent the 95% confidence interval, and the black line indicates the median N_e_ estimate.

## Discussion

This study uses phylogenomics to reconstruct the population history of the “harlequin frogs” (*Atelopus*) endemic to the Sierra Nevada de Santa Marta. All four morphospecies were recovered as robustly supported monophyletic lineages, corroborating the findings of Lötters et al., (2025) and Plewnia et al., (2026) using mitochondrial DNA. This phylogenetic pattern is congruent with clustering (PCA and DAPC) and ancestry (ADMIXTURE, fastSTRUCTURE, and fineRADstructure) analyses, all of which identified the species as distinct, highly structured genetic clusters.

Despite the clear boundaries between species, the phylogenetic relationships among them were discordant in our nuclear dataset. Nevertheless, our best species tree suggested by topology testing is congruent with the highly supported relationships (> 0.95) identified by Plewnia et al. (2026) using mitogenomics and short mitochondrial markers, which also matches the mitochondrial BI tree of Lötters et al. (2025). This topology resolves *A. laetissimus* and *A. arsyecue* as sister species, *A. carrikeri* as a sister to this clade, and finally, *A. nahumae* as a sister species to the other species.

Demographic modeling in GADMA yielded divergence times that are considerably younger than the estimates presented by Plewnia et al. (2026), who used BEAST calibrated with log-normal dating point from the oldest available bufonid fossil. The most narrow discrepancy between these studies occurs at the *A.arsyecue*-*A.carrikeri* node, which differs by ∼4.8 Ma. This discordance can be explained by the contrasting methods and data types employed. While BEAST estimates tend to overestimate divergence times by neglecting the coalescent process, leading to pronounced errors in the presence of large ancestral populations and heavy ILS (Angelis & Dos Reis, 2015) our analysis may suffer from a downward bias. Since our data is composed by short reads, the sequence length parameter required to estimate demography might be underestimated, thus also underestimating divergence times, GADMA2 also possesses inherent temporal limitations, losing accuracy for events deeper than 10N_anc_ generations ago (Noskova et al., 2022).

Another explanation for this discrepancy could be attributed to the nature of the data used in the analysis, since the mitochondrial genome is haploid and generally maternally inherited in most animals, its effective population size is fourthfold smaller than that of nuclear DNA (Hudson & Turelli, 2003; Zink & Barrowclough, 2008). Consequently, mtDNA normally completes lineage sorting at a faster rate, allowing ancestral polymorphisms to be lost faster than in the nuclear genome (D. J. Funk & Omland, 2003), although lower capacity to resolve species is also well known in some cases as consequence of different evolutionary processes acting on this molecule (e.g. *Ursus maritimus* and *Melinaea* and *Mechanitis* butterflies) (Lan et al., 2022; Van Der Heijden et al., 2025).

Moreover, mtDNA dating overestimation is exacerbated when calibration points are sparse and time spans are large (Hurley et al., 2007), as high mitochondrial mutation rates can cause phylogenetic models to inflate absolute split times (up to 3–10 times the real value) compared to nuclear datasets (Zheng et al., 2011). Studies have revealed that dating can be misleading with mtDNA if few calibration points are used, specifically if the lineage used is old and time span is large (Ho & Endicott, 2008; Zheng et al., 2011). Ultimately, resolving these competing biases and reconciling the disparity between types of data will likely require whole-genome sequencing (WGS) data.

Notably, there was a discrepancy in historical gene flow patterns between demography and introgression test, where demographic models suggest gene flow while ABBA-BABA based statistics showed no evidence of it. Smith & Hahn (2024) demonstrated that even when the data is consistent with an isolation-only (IM) model, different types of natural selection can cause an overestimation of migration. Furthermore, in the absence of selection, this overestimation can also occur in species with recent divergence times (< 1.5 Ma; Figure 4). On the other hand, statistics evaluating ABBA-BABA patterns tend to be biased toward detecting gene flow in its absence, driven by different phenomena (Frankel & Ané, 2023; Tournebize & Chikhi, 2024; Tricou et al., 2022), even so, these statistics display greater sensitivity in detecting gene flow in small populations (Zheng & Janke, 2018) and are robust to demographic shifts such as bottlenecks (De Martino et al., in prep.).

Given the limitations described for the two previous approaches, ABBA-BABA seems to be more conservative, thus the absence of gene flow among the *Atelopus* species of the SNSM is the most likely scenario. For this reason the observed pattern of discordance in phylogenetic relationships and shared ancestry among these species is likely driven by ILS. In fact, the estimated probability of this phenomenon for these toads is ∼9.08%, which is considered high in the literature (Blumer et al., 2025; Knief et al., 2024). Consequently, the confluence of recent divergence, historical isolation without gene flow and the presence of ILS, points toward a rapid speciation event within the SNSM *Atelopus* species.

Whether the speciation process between the four *Atelopus* species of the SNSM occurred along an ancient or recent timeline, the influence of ILS remains a highly robust explanation for the observed phylogenetic discordance. If the radiation is indeed recent (< 1.5 Ma; Figure 4), a high degree of ILS is expected simply due to the short evolutionary time elapsed since divergence, which inherently favors the retention of ancestral variation (Alaei Kakhki et al., 2023; Knief et al., 2024). However, the relevance of ILS is not restricted to recent radiations. Even if the divergence events took place much deeper in time, such that sufficient generations have passed for ancestral polymorphisms to become fully fixed within descendant lineages, the historical footprint of stochastic coalescent dynamics will still persist exclusively as discordance among the genealogies of different loci (Takahashi et al., 2001). This phenomenon has been widely documented across diverse animal radiations, such as in Lake Tanganyika cichlid fishes where ancient rapid splitting left a signature of gene tree conflict despite fixed polymorphisms (Takahashi et al., 2001), or more recently in Asian warty newts, where ILS and ancient hybridization collectively shaped a complex reticulate radiation (Tao et al., 2026). These systems demonstrate that when rapid lineage splitting coincides with large ancestral population sizes, ILS could generate long-lasting locus-specific phylogenetic conflict that can be detected long after divergence has occurred (Suh et al., 2015; Szöllősi et al., 2015).

Consistent with a scenario of no gene flow, absolute divergence values (D_xy_) between species pairs were higher than net divergence values (D_a_), possibly due to the retention of ancestral alleles that have segregated differentially, contributing to lineage differentiation (Cruickshank & Hahn, 2014; Pease et al., 2016; Yan et al., 2026). Furthermore, D_a_ values being slightly lower than D_xy_ instead of abruptly decreasing also reflects the presence of recent divergence that accumulated throughout speciation, implying that this process was influenced by both ancestral variation as well as independent differentiation.

The average net divergence (D_a_) values for each pair of species fell below the lower bound of the so-called “gray zone of speciation” although the D_a_ distribution spans several values both within and above this zone. Specifically, the number of outlier loci across pairwise comparisons was 5 for the *A. laetissimus*–*A. arsyecue*, 28 for *A. laetissimus*–*A. carrikeri*, 39 for *A. arsyecue*–*A. carrikeri*, 63 for *A. arsyecue*–*A. nahumae*, 64 for *A. nahumae*–*A. carrikeri*, and 69 for *A. laetissimus*–*A. nahumae*. Having ruled out gene flow, the importance of these outliers in speciation with ILS remains unknown and could instead be accumulated after speciation.

Although it has been suggested that physical barriers alone do not contribute substantially to IR (Westram et al., 2022), the lack of gene flow between morphospecies may be partly explained by the specific geographical and ecological conditions of the SNSM. This mountain range features high environmental heterogeneity as well as topographic complexity (Hatzenbühler et al., 2022; Montes et al., 2010; Strewe & Navarro, 2021), which may restrict connectivity and dispersal among populations of these species. However, it was impossible to sample the full distribution of each species in order to assess the importance of these topographical and climatic factors in promoting genetic isolation due to the region’s historic and ongoing armed conflict, which severely limits field accessibility in certain areas of the SNSM. In fact, geography as a barrier to interspecies gene flow has been reported in other *Atelopus* species found in the Guiana Shield, as well as *A. manauensis* and *A. varius*, distributed in northeastern Brazil and western Panama, respectively (Jorge et al., 2022; Noonan & Gaucher, 2005; Richards-Zawacki, 2009).

Ecological and behavioral differences can act as reproductive barriers between species that come into geographical contact and thus facilitate or maintain speciation. For example, microhabitat preference in the case of *A. laetissimus* who inhabits humid mid-elevation forests with dense vegetation and low light levels, and exhibits primarily nocturnal activity, whereas *A. nahumae* occupies more open forests and has shown mostly diurnal behavior (Granda-Rodríguez et al., 2020; Rueda Solano et al., 2016). Although these two species come into contact by sharing breeding areas near water sources and a single interspecific amplexus recorded (J. L. Pérez-Gonzalez et al., 2020), however, the result of this amplexus and whether or not it was successful was not recorded (i.e. no data exist about postzygotic isolation between these species). Given that there has only been 1 singular event and no natural hybrids documented between these species, differences in microhabitat preferences and activity periods may have contributed to keep these species isolated as was demonstrated in other animals such as tree frogs from the *Hyla* genus and *Heliconius* butterflies from the *melpomene–cydno* complex (Borzée et al., 2016; Montgomery et al., 2021). The effect of different types of selection in reproductive isolation such as strong intrasexual competition among males as well as vocalizations of males and females, which have been documented in *A. laetissimus*, also occur in the other species and whether they could contribute to reproductive isolation (RI) remains unknown (Rueda-Solano et al., 2020, 2022). Therefore, further integration of whole-genome sequencing (WGS) and behavioral studies is needed to identify the diverse evolutionary phenomena that have shaped divergence within this group of toads.

The specific scenario of rapid speciation could also be driven by changes in the genomic architecture that halters gene flow. For instance, ploidy shifts are known drivers of speciation in plants (Wood et al., 2009) and certain fish lineages (Rennison et al., 2012; Venkatesh, 2003), regulatory incompatibilities involving sex chromosomes acting as rapid triggers for hybrid sterility seen in birds and mice (Delph & Demuth, 2016; Irwin, 2018; Johnson & Lachance, 2012; Larson et al., 2016) or copy number variations (CNVs) in genes encoding ionotropic receptors, potentially linked to ecological adaptations in the genus *Heliconius* (Van Schooten et al., 2016). Alternatively, an increased effect of genetic drift caused by small population sizes, geographic isolation, and the occurrence of repeated bottlenecks was hypothesized to facilitate rapid speciation events in vertebrate fish (Black et al., 2024) as well as mammals (W. C. Funk et al., 2016; Kalirad et al., 2024; Qi et al., 2025). This evolutionary scenario could be mirrored in the *Atelopus* species of the SNSM since our analyses evidence an overall contraction in population size, high environmental discontinuity, and apparent recent speciation, lack of genetic connectivity, high heterogeneity in vegetation cover and complex topography along this isolated mountain peaks. Nevertheless, the specific mechanisms underlying the disruption of gene flow among SNSM species remain unidentified, including whether they involve pre- or postzygotic barriers to reproductive isolation. Establishing the role of genetic drift, natural selection, sexual selection or a combination of them in the speciation of these toads requires additional studies.

The historical fluctuation in population size for these species could be attributed to repeated glacial periods starting 200.000 years ago until the Last Glacial Maximum between 21–18 ka (Lachniet & Vazquez-Selem, 2005; López-Moreno et al., 2020; Shin et al., 2020). However, glaciations and their effects on the SNSM have not been studied in depth (Cornelius Raasveldt, 2017; Lachniet & Vazquez-Selem, 2005; Schubert & Clapperton, 1990), so demographic changes in these species may not be strictly synchronized with climatic pulses and may have been modulated by other factors.

From a conservation perspective, the estimated effective population size in *A.laetissimus* (N_e_ = 342) surpasses the lower threshold of the 50/500 rule (Clarke et al., 2024; Franklin et al., 1980; Jamieson & Allendorf, 2012), suggesting that this species still possesses sufficient genetic diversity to limit the effects of inbreeding in the short term. However, some studies suggest that these thresholds are not applicable to all taxa and must be adjusted to the life history traits of each species; therefore, they have been proposed to be increased to N_e_ 100/1000 rule (Franklin et al., 2014; Pérez-Pereira et al., 2022; Waples, 2025). With regard to this modification, the estimated N_e_ value for *A. laetissimus* would be even further from the upper threshold, considered optimal for maintaining a population’s adaptive capacity. However, there are no population census records for the other species, so it is not possible to assess their status under this conservation rule.

## Conclusions

In conclusion, our phylogenomic analyses robustly resolve four independent, monophyletic lineages within the *Atelopus* present in the SNSM, which along with genetic structure, supports their status as distinct species. The absence of contemporary gene flow, combined with the likely presence of incomplete lineage sorting (ILS), supports a scenario of rapid speciation, where an ancestral lineage split rapidly forming new genetically isolated species. While our demographic models point to recent diversification timeline, whole genome sequencing (WGS) will be essential to definitively reconcile divergence dates across nuclear and mitochondrial datasets. Crucially, the endemic nature and lack of genetic connectivity across these isolated mountain peaks may significantly exacerbate the vulnerability of these species to global warming and emerging diseases, with *A.laestissimus* facing particularly high risk of future inbreeding.

## Acknowledgements

The collection of amphibians was authorized by Autoridad Nacional de Licencias Ambientales (ANLA) through the permit granted to Fundación Atelopus (Resolución 1441 de 2020). The specimens were treated according to the ethics committee (number CV1599) of Universidad del Rosario. We are grateful to Fundación Atelopus for their assistance in collecting the anuran specimens in Sierra Nevada de Santa Marta (SNSM). We thank Noskova et al. (2022), maintainer of GADMA2, for her insightful help with the implementation of this software.

## Funding

This research was supported by the joint funding call between Universidad del Rosario and Universidad del Magdalena (*Convocatoria de Proyectos 2022-1*).

## Notes

### Competing Interest Statement

The authors have declared no competing interest.

