## Supplementary material for "Rapid speciation in Harlequin Toads (*Anura*: *Bufonidae*) endemic to the Sierra Nevada de Santa Marta"

**Table S1.** Individuals sampled by morphospecies, along with their collection sites.

| Especie | Código | Departamento | Municipio | Localidad | Latitud | Longitud |
|---|---|---|---|---|---|---|
| *A. carrikeri* | 18 | Magdalena | Ciénaga | Páramo de Cebolletas | 10.90103 | -73.917913 |
| *A. carrikeri* | 39 | Magdalena | Ciénaga | Páramo de Cebolletas | 10.90103 | -73.917913 |
| *A. carrikeri* | 23 | Magdalena | Ciénaga | Páramo de Cebolletas | 10.90103 | -73.917913 |
| *A. carrikeri* | 79 | Magdalena | Ciénaga | Páramo de Cebolletas | 10.90103 | -73.917913 |
| *A. carrikeri* | 81 | Magdalena | Ciénaga | Páramo de Cebolletas | 10.90103 | -73.917913 |
| *A. carrikeri* | 45 | Magdalena | Ciénaga | Páramo de Cebolletas | 10.90103 | -73.917913 |
| *A. carrikeri* | 76 | Magdalena | Ciénaga | Páramo de Cebolletas | 10.90103 | -73.917913 |
| *A. carrikeri* | 36 | Magdalena | Ciénaga | Páramo de Cebolletas | 10.90103 | -73.917913 |
| *A. carrikeri* | 20 | Magdalena | Ciénaga | Páramo de Cebolletas | 10.90103 | -73.917913 |
| *A. carrikeri* | 35 | Magdalena | Ciénaga | Páramo de Cebolletas | 10.90103 | -73.917913 |
| *A. carrikeri* | 19 | Magdalena | Ciénaga | Páramo de Cebolletas | 10.90103 | -73.917913 |
| *A. carrikeri* | 85 | Magdalena | Ciénaga | Páramo de Cebolletas | 10.90103 | -73.917913 |
| *A. carrikeri* | 84 | Magdalena | Ciénaga | Páramo de Cebolletas | 10.90103 | -73.917913 |

| | | | | | | |
|---|---|---|---|---|---|---|
| *A. carrikeri* | 37 | Magdalena | Ciénaga | Páramo de Cebolletas | 10.90103 | -73.917913 |
| *A. carrikeri* | 33 | Magdalena | Ciénaga | Páramo de Cebolletas | 10.90103 | -73.917913 |
| *A. nahumae* | 17 | Magdalena | Santa Marta | San Lorenzo | 11.12580556 | -74.0545556 |
| *A. nahumae* | 15 | Magdalena | Santa Marta | San Lorenzo | 11.12580556 | -74.0545556 |
| *A. nahumae* | 1 | Magdalena | Santa Marta | San Lorenzo | 11.12580556 | -74.0545556 |
| *A. nahumae* | 77 | Magdalena | Santa Marta | San Lorenzo | 11.12580556 | -74.0545556 |
| *A. nahumae* | 27 | Magdalena | Ciénaga | San Pedro de la Sierra | 10.897983 | -73.9934 |
| *A. nahumae* | 32 | Magdalena | Ciénaga | San Pedro de la Sierra | 10.897983 | -73.9934 |
| *A. nahumae* | 46 | Magdalena | Ciénaga | San Pedro de la Sierra | 10.897983 | -73.9934 |
| *A. nahumae* | 90 | Magdalena | Santa Marta | San Lorenzo | 11.12580556 | -74.0545556 |
| *A. nahumae* | 47 | Magdalena | Santa Marta | San Lorenzo | 11.12580556 | -74.0545556 |
| *A. nahumae* | 66 | Magdalena | Santa Marta | San Lorenzo | 11.12580556 | -74.0545556 |
| *A. nahumae* | 48 | Magdalena | Santa Marta | San Lorenzo | 11.12580556 | -74.0545556 |
| *A. nahumae* | 68 | Magdalena | Santa Marta | San Lorenzo | 11.12580556 | -74.0545556 |
| *A. nahumae* | 87 | Magdalena | Santa Marta | San Lorenzo | 11.12580556 | -74.0545556 |
| *A. nahumae* | 70 | Magdalena | Santa Marta | San Lorenzo | 11.12580556 | -74.0545556 |
| *A. nahumae* | 72 | Magdalena | Santa Marta | San Lorenzo | 11.12580556 | -74.0545556 |
| *A. nahumae* | 67 | Magdalena | Santa Marta | San Lorenzo | 11.12580556 | -74.0545556 |
| *A. nahumae* | 61 | Magdalena | Santa Marta | San Lorenzo | 11.12580556 | -74.0545556 |

| | | | | | | |
|---|---|---|---|---|---|---|
| *A. nahumae* | 73 | Magdalena | Santa Marta | San Lorenzo | 11.12580556 | -74.0545556 |
| *A. nahumae* | 59 | Magdalena | Santa Marta | San Lorenzo | 11.12580556 | -74.0545556 |
| *A. nahumae* | 88 | Magdalena | Santa Marta | San Lorenzo | 11.12580556 | -74.0545556 |
| *A. nahumae* | 89 | Magdalena | Santa Marta | San Lorenzo | 11.12580556 | -74.0545556 |
| *A. nahumae* | 69 | Magdalena | Santa Marta | San Lorenzo | 11.12580556 | -74.0545556 |
| *A. arsyecue* | 94 | Cesar | Valledupar | Sogrome | 10.7333 | -73.48333 |
| *A. arsyecue* | 55 | Cesar | Valledupar | Sogrome | 10.7333 | -73.48333 |
| *A. arsyecue* | 52 | Cesar | Valledupar | Sogrome | 10.7333 | -73.48333 |
| *A. arsyecue* | 64 | Cesar | Valledupar | Sogrome | 10.7333 | -73.48333 |
| *A. arsyecue* | 63 | Cesar | Valledupar | Sogrome | 10.7333 | -73.48333 |
| *A. arsyecue* | 62 | Cesar | Valledupar | Sogrome | 10.7333 | -73.48333 |
| *A. arsyecue* | 97 | Cesar | Valledupar | Sogrome | 10.7333 | -73.48333 |
| *A. arsyecue* | 96 | Cesar | Valledupar | Sogrome | 10.7333 | -73.48333 |
| *A. arsyecue* | 82 | Cesar | Valledupar | Sogrome | 10.7333 | -73.48333 |
| *A. arsyecue* | 95 | Cesar | Valledupar | Sogrome | 10.7333 | -73.48333 |
| *A. arsyecue* | 65 | Cesar | Valledupar | Sogrome | 10.7333 | -73.48333 |
| *A. arsyecue* | 54 | Cesar | Valledupar | Sogrome | 10.7333 | -73.48333 |
| *A. arsyecue* | 50 | Cesar | Valledupar | Sogrome | 10.7333 | -73.48333 |
| *A. arsyecue* | 51 | Cesar | Valledupar | Sogrome | 10.7333 | -73.48333 |

| | | | | | | |
|---|---|---|---|---|---|---|
| *A. arsyecue* | 53 | Cesar | Valledupar | Sogrome | 10.7333 | -73.48333 |
| *A. laetissimus* | 14 | Magdalena | Santa Marta | San Lorenzo | 11.12580556 | -74.0545556 |
| *A. laetissimus* | 80 | Magdalena | Santa Marta | San Lorenzo | 11.12580556 | -74.0545556 |
| *A. laetissimus* | 41 | Magdalena | Santa Marta | San Lorenzo | 11.12580556 | -74.0545556 |
| *A. laetissimus* | 49 | Magdalena | Santa Marta | San Lorenzo | 11.12580556 | -74.0545556 |
| *A. laetissimus* | 16 | Magdalena | Santa Marta | San Lorenzo | 11.12580556 | -74.0545556 |
| *A. laetissimus* | 26 | Magdalena | Santa Marta | San Lorenzo | 11.12580556 | -74.0545556 |
| *A. laetissimus* | 9 | Magdalena | Santa Marta | San Lorenzo | 11.12580556 | -74.0545556 |
| *A. laetissimus* | 78 | Magdalena | Santa Marta | San Lorenzo | 11.12580556 | -74.0545556 |
| *A. laetissimus* | 5 | Magdalena | Santa Marta | San Lorenzo | 11.12580556 | -74.0545556 |
| *A. laetissimus* | 4 | Magdalena | Santa Marta | San Lorenzo | 11.12580556 | -74.0545556 |
| *A. laetissimus* | 21 | Magdalena | Santa Marta | San Lorenzo | 11.12580556 | -74.0545556 |
| *A. laetissimus* | 22 | Magdalena | Santa Marta | San Lorenzo | 11.12580556 | -74.0545556 |
| *A. laetissimus* | 40 | Magdalena | Ciénaga | San Pedro de la Sierra | 10.897983 | -73.9934 |
| *A. laetissimus* | 42 | Magdalena | Ciénaga | San Pedro de la Sierra | 10.897983 | -73.9934 |
| *A. laetissimus* | 34 | Magdalena | Ciénaga | San Pedro de la Sierra | 10.897983 | -73.9934 |
| *A. laetissimus* | 8 | Magdalena | Ciénaga | San Pedro de la Sierra | 10.897983 | -73.9934 |
| *A. laetissimus* | 25 | Magdalena | Ciénaga | San Pedro de la Sierra | 10.897983 | -73.9934 |
| *A. laetissimus* | 38 | Magdalena | Ciénaga | San Pedro de la Sierra | 10.897983 | -73.9934 |

| | | | | | | |
|---|---|---|---|---|---|---|
| *A. laetissimus* | 28 | Magdalena | Ciénaga | San Pedro de la Sierra | 10.897983 | -73.9934 |
| *A. laetissimus* | 57 | Magdalena | Santa Marta | San Lorenzo | 11.12580556 | -74.0545556 |
| *A. laetissimus* | 58 | Magdalena | Santa Marta | San Lorenzo | 11.12580556 | -74.0545556 |
| *A. laetissimus* | 56 | Magdalena | Santa Marta | San Lorenzo | 11.12580556 | -74.0545556 |
| *A. laetissimus* | 86 | Magdalena | Santa Marta | San Lorenzo | 11.12580556 | -74.0545556 |
| *A. laetissimus* | 92 | Magdalena | Santa Marta | San Lorenzo | 11.12580556 | -74.0545556 |
| *A. laetissimus* | 91 | Magdalena | Santa Marta | San Lorenzo | 11.12580556 | -74.0545556 |
| *A. laetissimus* | 93 | Magdalena | Santa Marta | San Lorenzo | 11.12580556 | -74.0545556 |
| *A. laetissimus* | 83 | Magdalena | Santa Marta | San Lorenzo | 11.12580556 | -74.0545556 |
| *A. laetissimus* | 71 | Magdalena | Santa Marta | San Lorenzo | 11.12580556 | -74.0545556 |
| *A. laetissimus* | 7 | La Guajira | Dibulla | Rio ancho | 11.11584 | -73.5513 |
| *A. laetissimus* | 3 | La Guajira | Dibulla | Rio ancho | 11.11584 | -73.5513 |
| *A. laetissimus* | 12 | La Guajira | Dibulla | Rio ancho | 11.11584 | -73.5513 |
| *A. laetissimus* | 74 | La Guajira | Dibulla | Rio ancho | 11.11584 | -73.5513 |
| *A. laetissimus* | 75 | La Guajira | Dibulla | Rio ancho | 11.11584 | -73.5513 |
| *A. laetissimus* | 2 | La Guajira | Dibulla | Rio ancho | 11.11584 | -73.5513 |
| *A. laetissimus* | 30 | La Guajira | Dibulla | Rio ancho | 11.11584 | -73.5513 |
| *A. laetissimus* | 24 | La Guajira | Dibulla | Rio ancho | 11.11584 | -73.5513 |
| *A. laetissimus* | 11 | La Guajira | Dibulla | Rio ancho | 11.11584 | -73.5513 |

| | | | | | | |
|---|---|---|---|---|---|---|
| *A.spurelli* | 6 | Chocó | Nuquí | El Amargal | 5.5705 | -77.5026 |
| *A.spurelli* | 44 | Chocó | Nuquí | El Amargal | 5.5705 | -77.5026 |
| *A. fronterizo* | 29 | Chocó | Acandí | Capurganá | 8.642550 | -77.395544 |
| *A. fronterizo* | 31 | Chocó | Acandí | Capurganá | 8.642551 | -77.395545 |
| *A. fronterizo* | 13 | Choco | Colombia | Acandí | 8.642552 | -77.395546 |

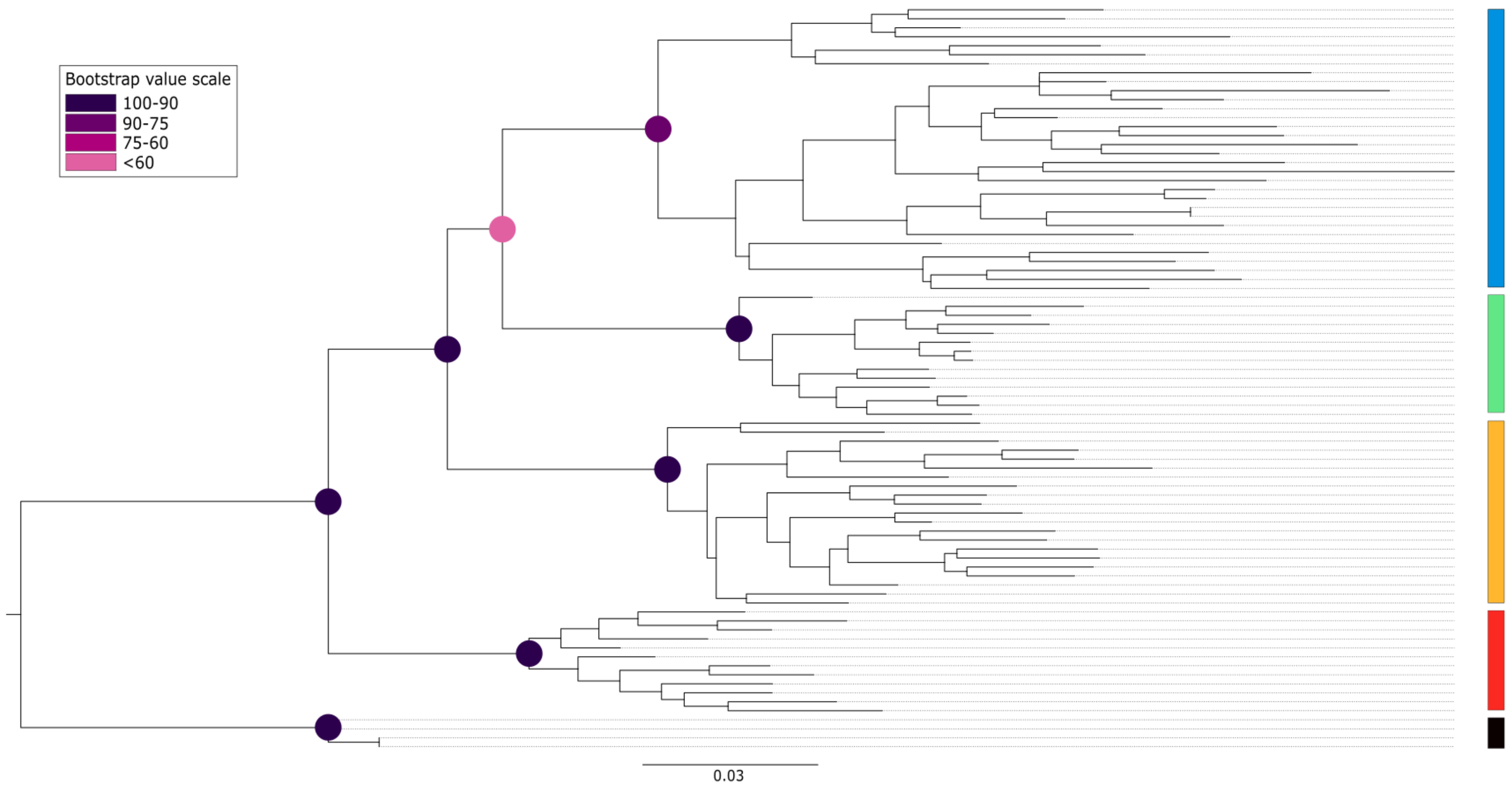


**Figure S1.** Maximum likelihood phylogenetic tree inferred with RAxML. Colored circles at the nodes indicate node support values for bootstrap ranging from purple (100% support) to pink (< 60% support). Species are identified by color: Blue: *A. laetissimus*, green: *A. arsyecue*, orange: *A. nahumae*, red: *A. carrikeri*, and black: Outgroup.

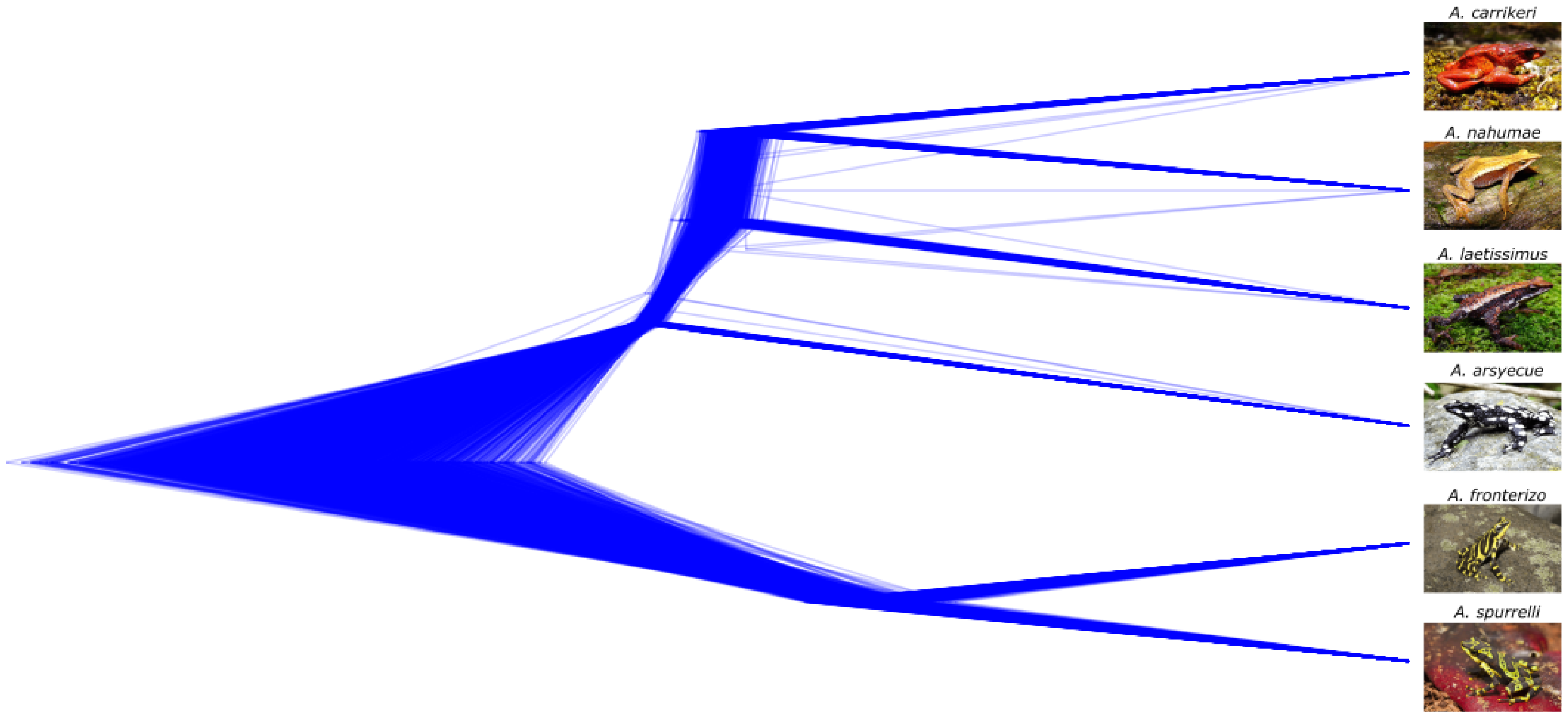


**Figure S2.** Density tree inferred using SNAPPER. Each line represents a topology (3,000 in total). Branch lengths were transformed using a pseudo-logarithmic scale to facilitate the visualization of short branches.

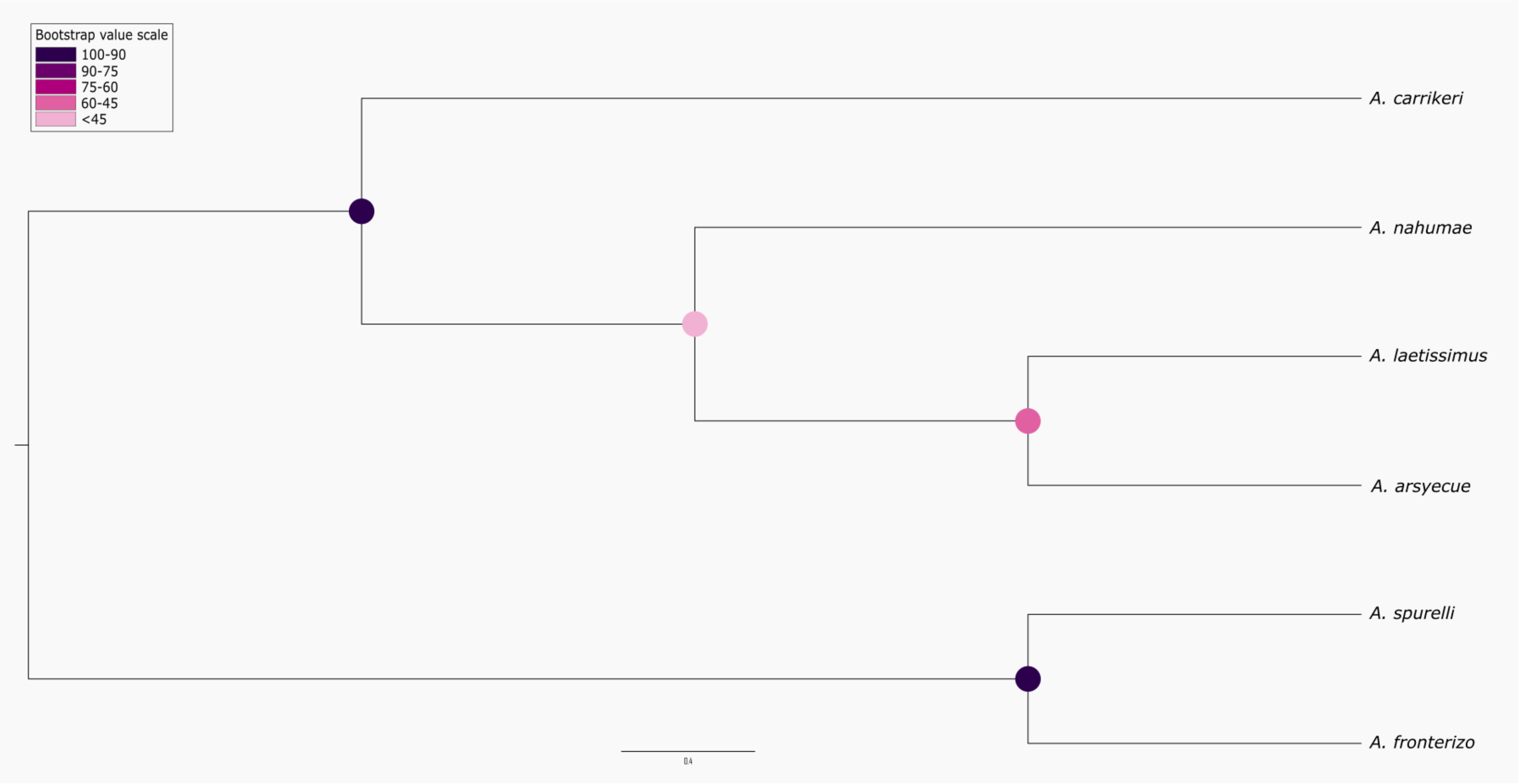


**Figure S3.** Multispecies coalescent tree inferred with SVDQuartets. Colored circles at the nodes indicate node support values for bootstrap ranging from purple (100% support) to pink (< 60% support).

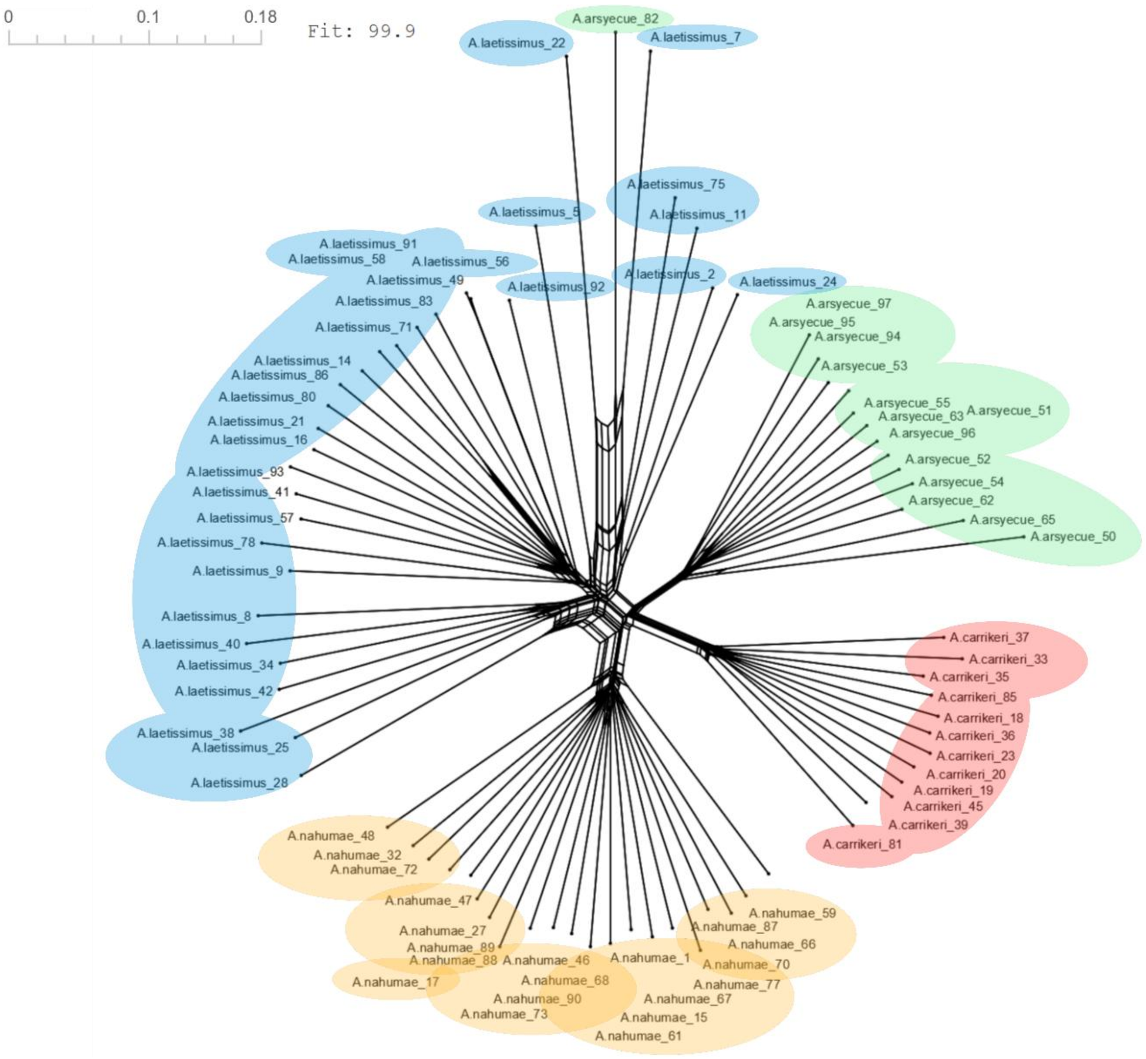


**Figure S4.** Neighbornet network. Species are identified by color: Blue: *A. laetissimus*, green: *A. arsyecue*, orange: *A. nahumae* and red: *A. carrikeri*.

**Table S2.** Log-likelihood values and Approximately Unbiased (AU) test results for the evaluated tree topologies. Statistical significance for the Approximately Unbiased (AU) test and the complementary tests: Kishino–Hasegawa (KH), Shimodaira–Hasegawa (SH), Expected Likelihood Weight (ELW), and bootstrap Proportion (BP), calculated using the RELL resampling procedure with 10,000 replicates.

| Topología de árbol | Método de inferencia (software) | logL | deltaL | bp-RELL | p-KH | p-SH | c-ELW | p-AU |
|---|---|---|---|---|---|---|---|---|
| ((*A. laetissimus*,*A. arsyecue*),*A. carrikeri*),*A. nahumae*) | Máxima verosimilitud (IQ-TREE 2) | -92164.3028 | 0 | 1 (+) | 1 (+) | 1 (+) | 1 (+) | 1 |
| ((*A. laetissimus*,*A. arsyecue*),*A. nahumae*),*A. carrikeri*) | Máxima verosimilitud (RAxML) / modelo coalescente multiespecies (SVDQuartest) | -93071.75509 | 907.45 | 0 (-) | 0 (-) | 0 (-) | 2.98E-76 (-) | 1.96E-33 |
| (((*A. carrikeri*,*A. nahumae*),*A. laetissimus*),*A. arsyecue*) | Modelo coalescente multiespecies (SNAPPER) | -92684.36247 | 520.06 | 0 (-) | 0 (-) | 0.007 (-) | 9.95E-113 (-) | 1.24E-06 |

deltaL : logL difference from the maximal logl in the set; bp-RELL : bootstrap proportion using RELL method (Kishino et al. 1990); p-KH : p-value of one sided Kishino-Hasegawa test (1989); p-SH : p-value of Shimodaira-Hasegawa test (2000); c-ELW : Expected Likelihood Weight (Strimmer & Rambaut 2002) y p-AU : p-value of approximately unbiased (AU) test (Shimodaira, 2002).

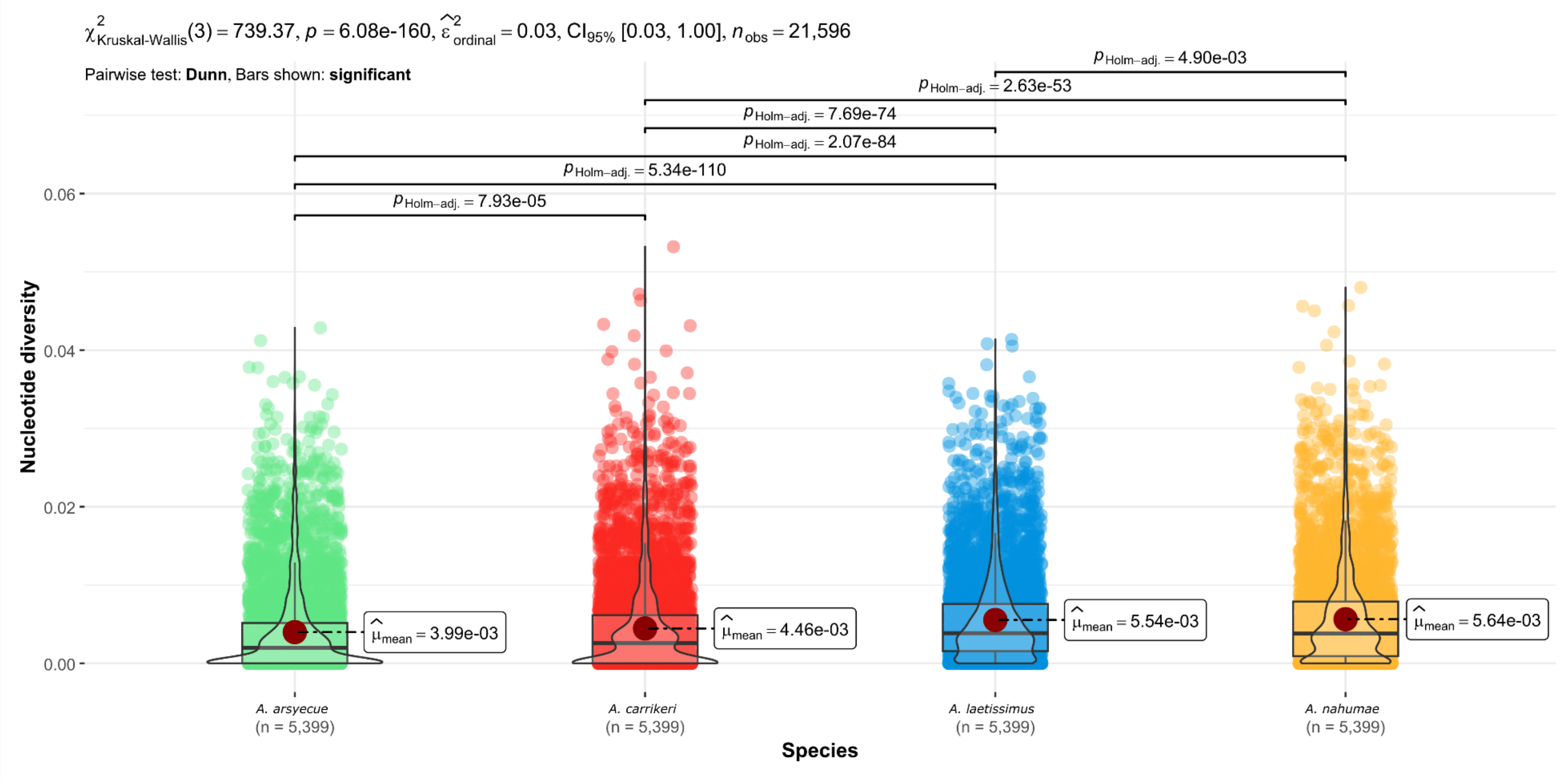


**Figure S5.** Nucleotide diversity per tag per morphospecies. The red circles indicate the mean, and the bars above show the statistical significance of the differences. Species are identified by color: Blue: *A. laetissimus*, green: *A. arsyecue*, orange: *A. nahumae* and red: *A. carrikeri*.

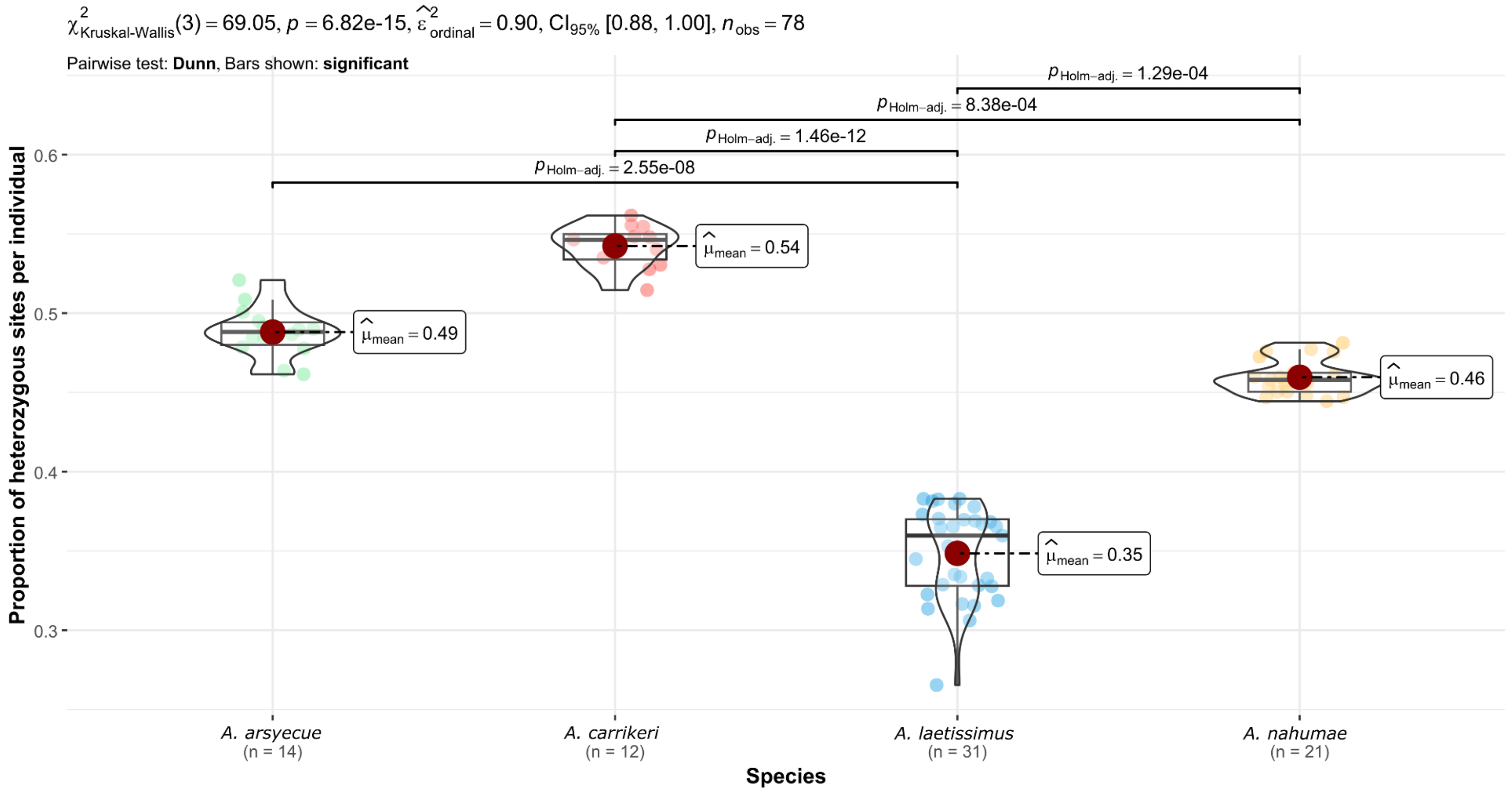


**Figure S6.** Proportion of heterozygosity per locus per individual per species. The red circles indicate the mean, and the bars above show the statistical significance of the differences. Species are identified by color: Blue: *A. laetissimus*, green: *A. arsyecue*, orange: *A. nahumae* and red: *A. carrikeri*.

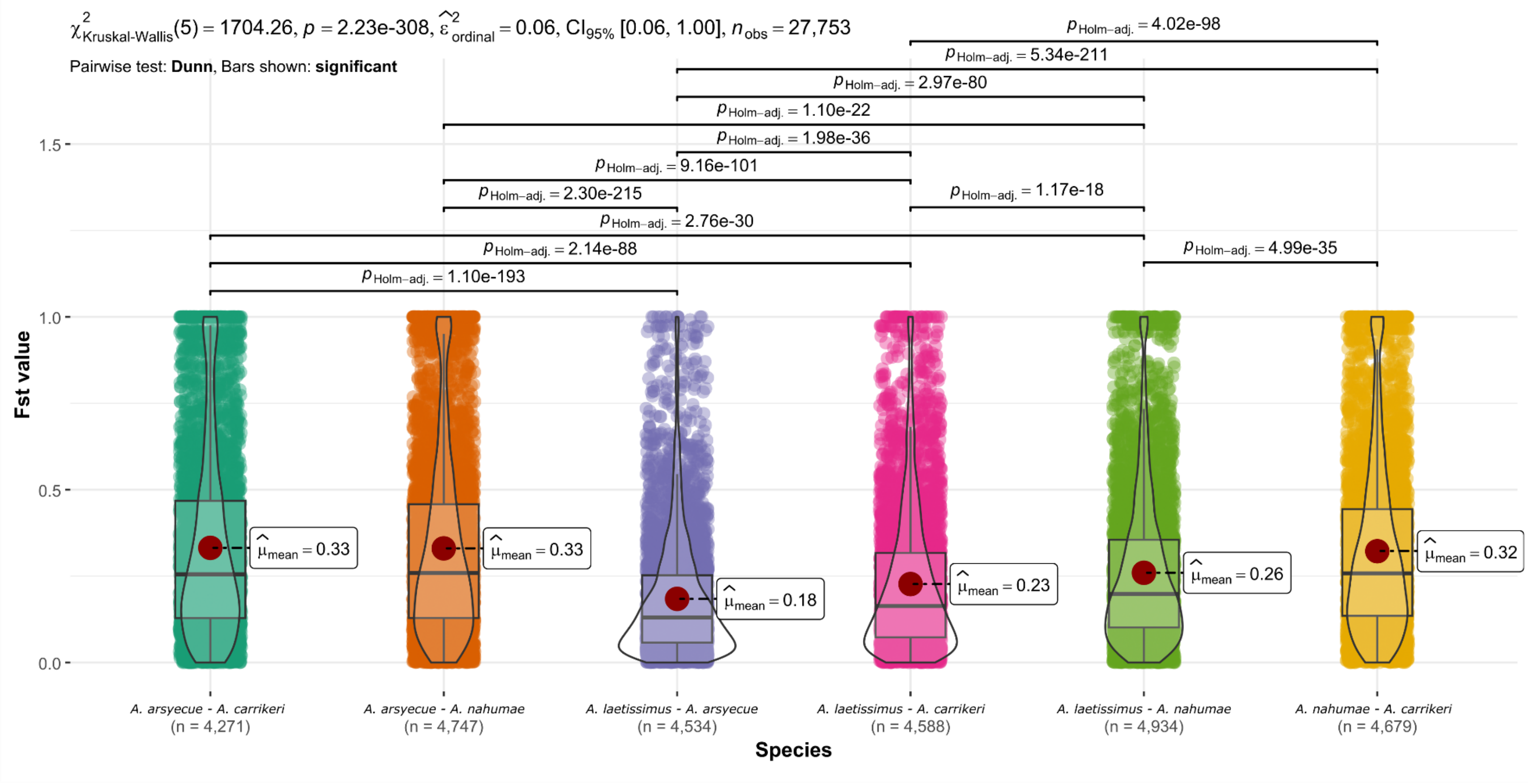


**Figure S7.** Relative genetic divergence ($F_{st}$) among species. Values for six pairwise comparisons of the four species. The points represent the analyzed tags, the red circles indicate the mean, the black horizontal line indicates the median, and the upper bars show the statistical significance of the differences.

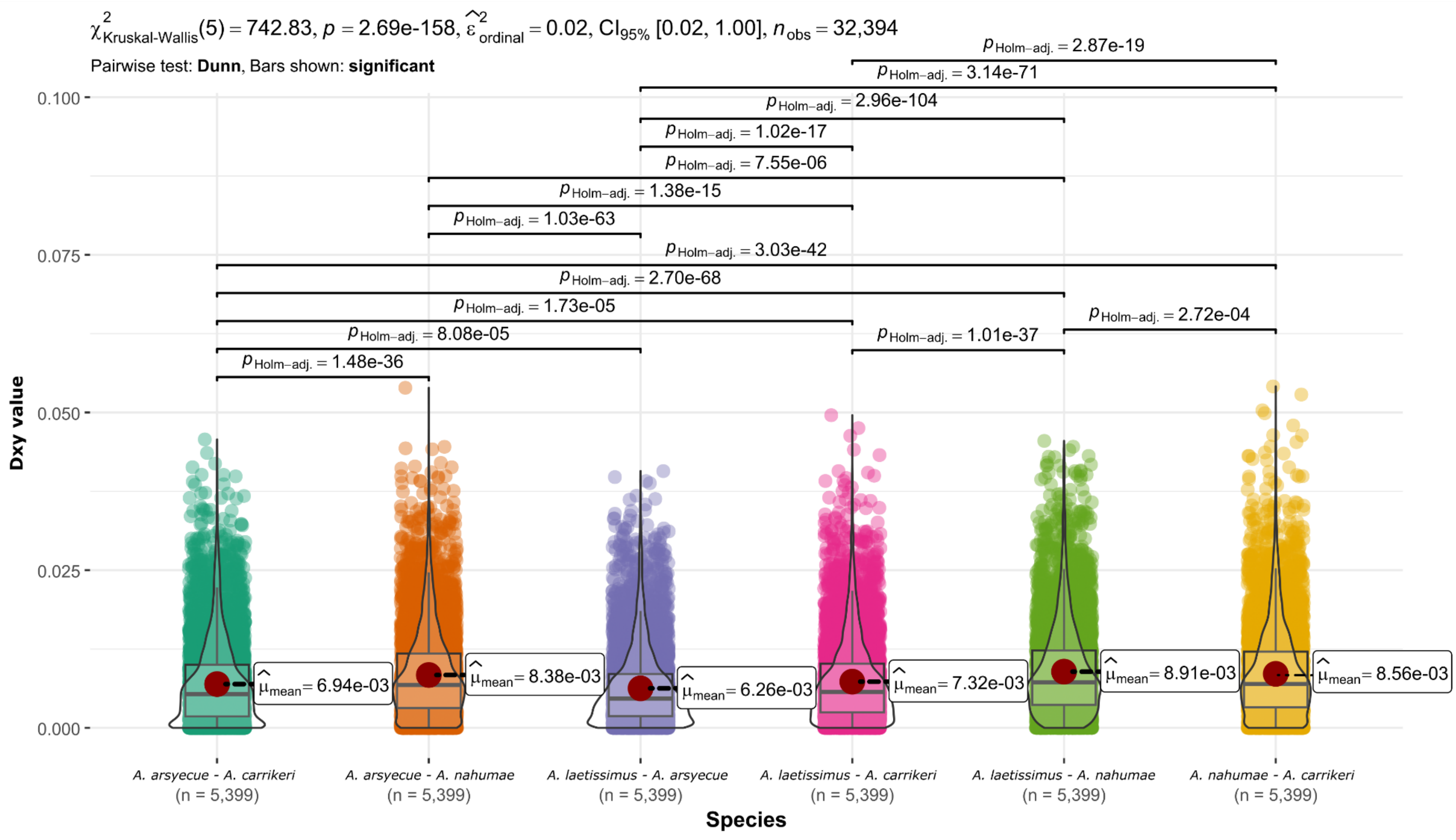


**Figure S8.** Absolute genetic divergence ($D_{xy}$) among species. Values for six pairwise comparisons of the four species. The dots represent the analyzed tags, the red circles indicate the mean, the black horizontal line indicates the median, and the upper bars show the statistical significance of the differences.

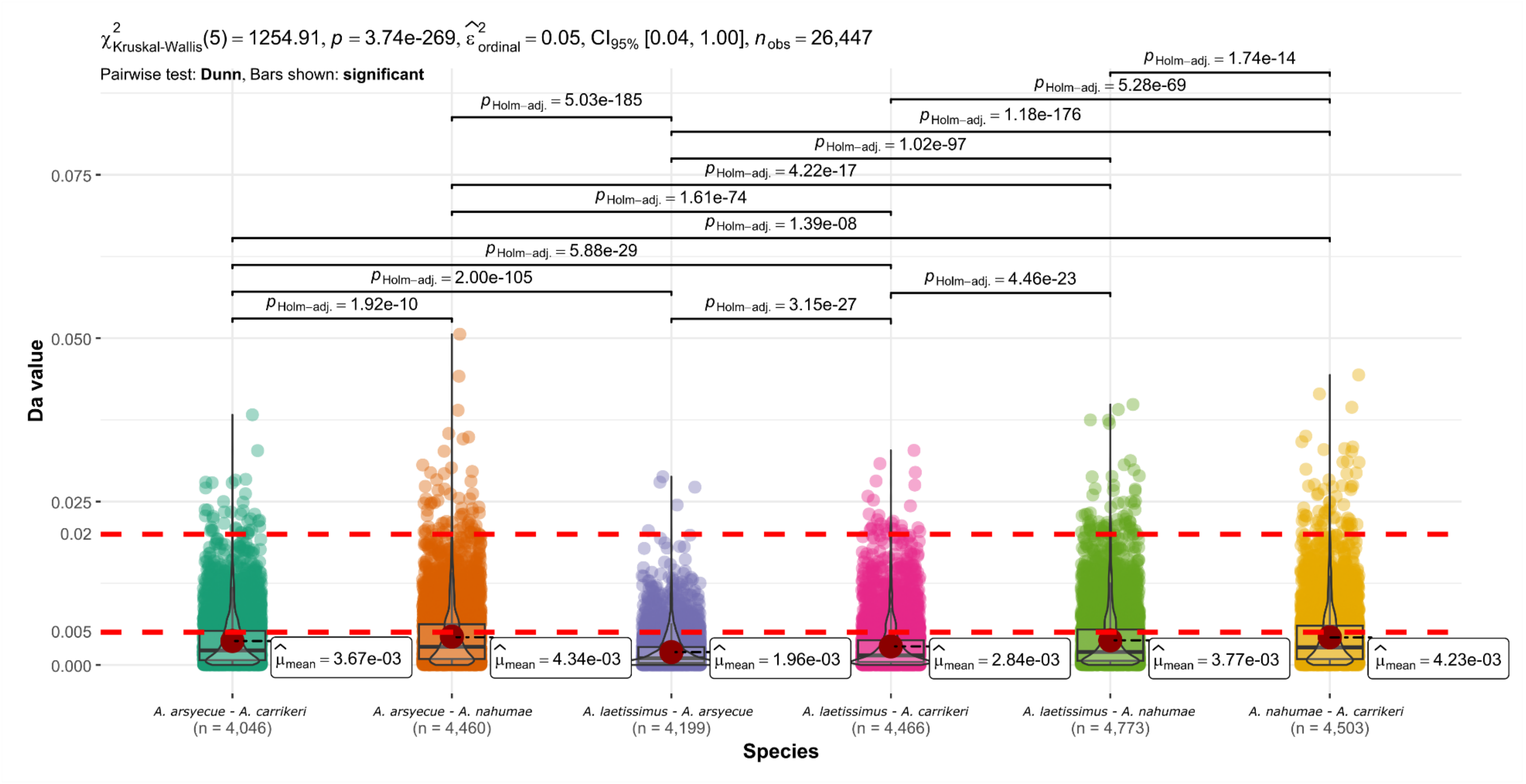


**Figure S9.** Net genetic divergence ($D_a$) among species. Values for six pairwise comparisons of the four species. The dots represent the analyzed tags, the red circles indicate the mean, the black horizontal line indicates the median, and the upper bars show the statistical significance of the differences. The boundaries of the "speciation gray zone" ($0.005 < D_a < 0.02$) are represented by the red horizontal lines.

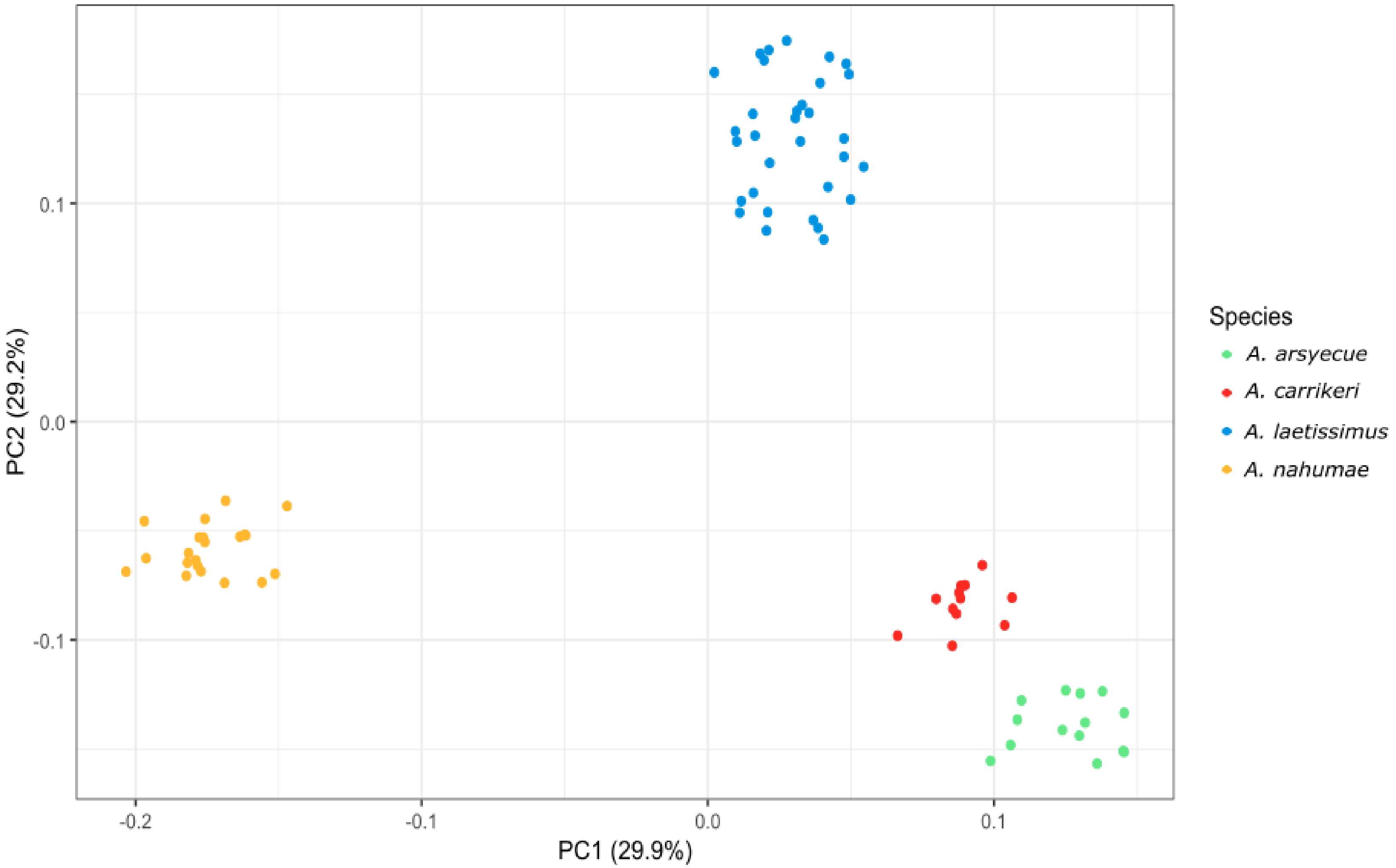


**Figure S10.** Principal component analysis (PCA) of genetic diversity among the species. Each point represents an individual. Species are identified by color: Blue: *A. laetissimus*, green: *A. arsyecue*, orange: *A. nahumae* and red: *A. carrikeri*.

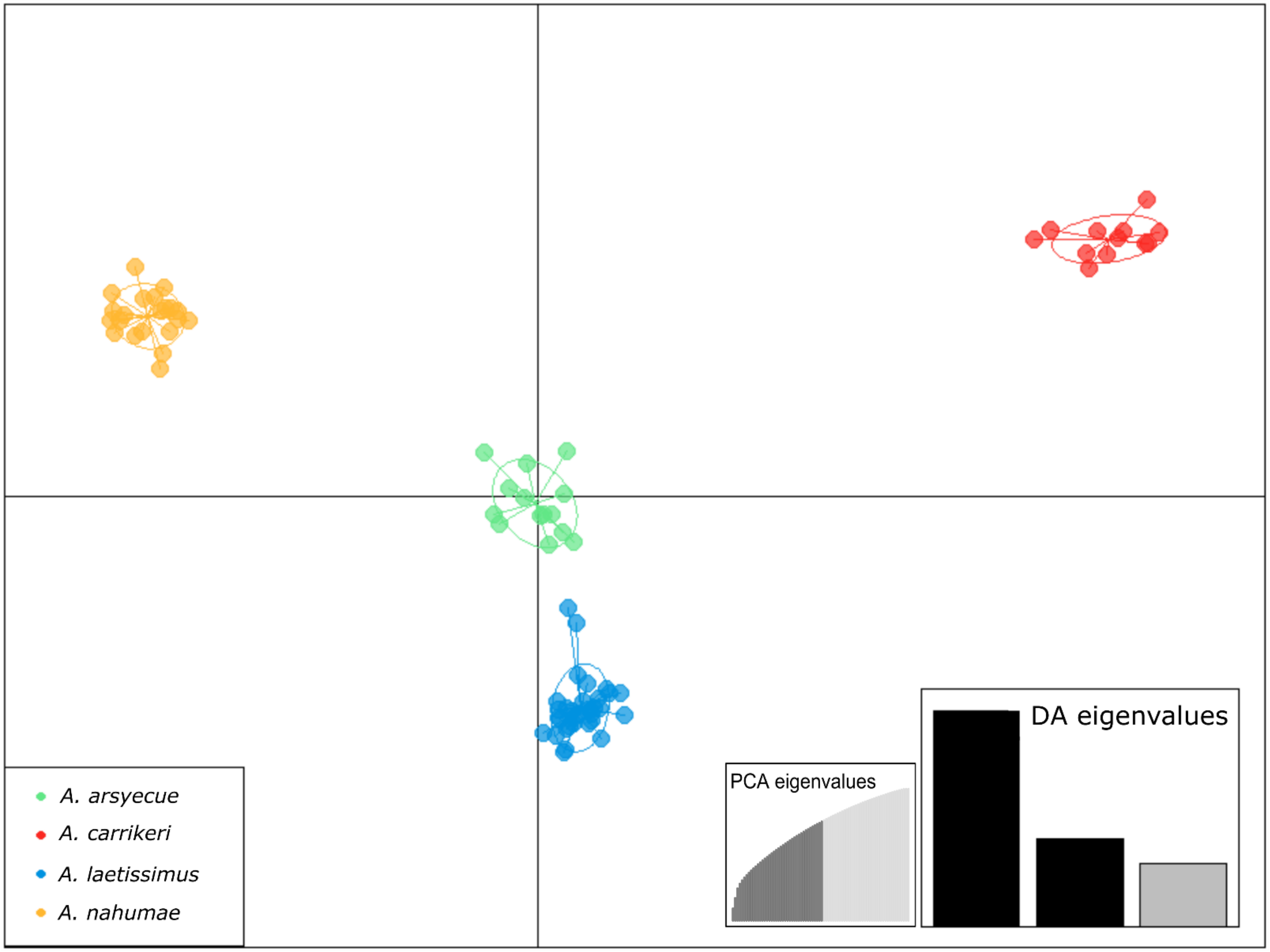


**Figure S11.** Discriminant Analysis of Principal Components (DAPC). Each point represents an individual. Species are identified by color: Blue: *A. laetissimus*, green: *A. arsyecue*, orange: *A. nahumae* and red: *A. carrikeri*.

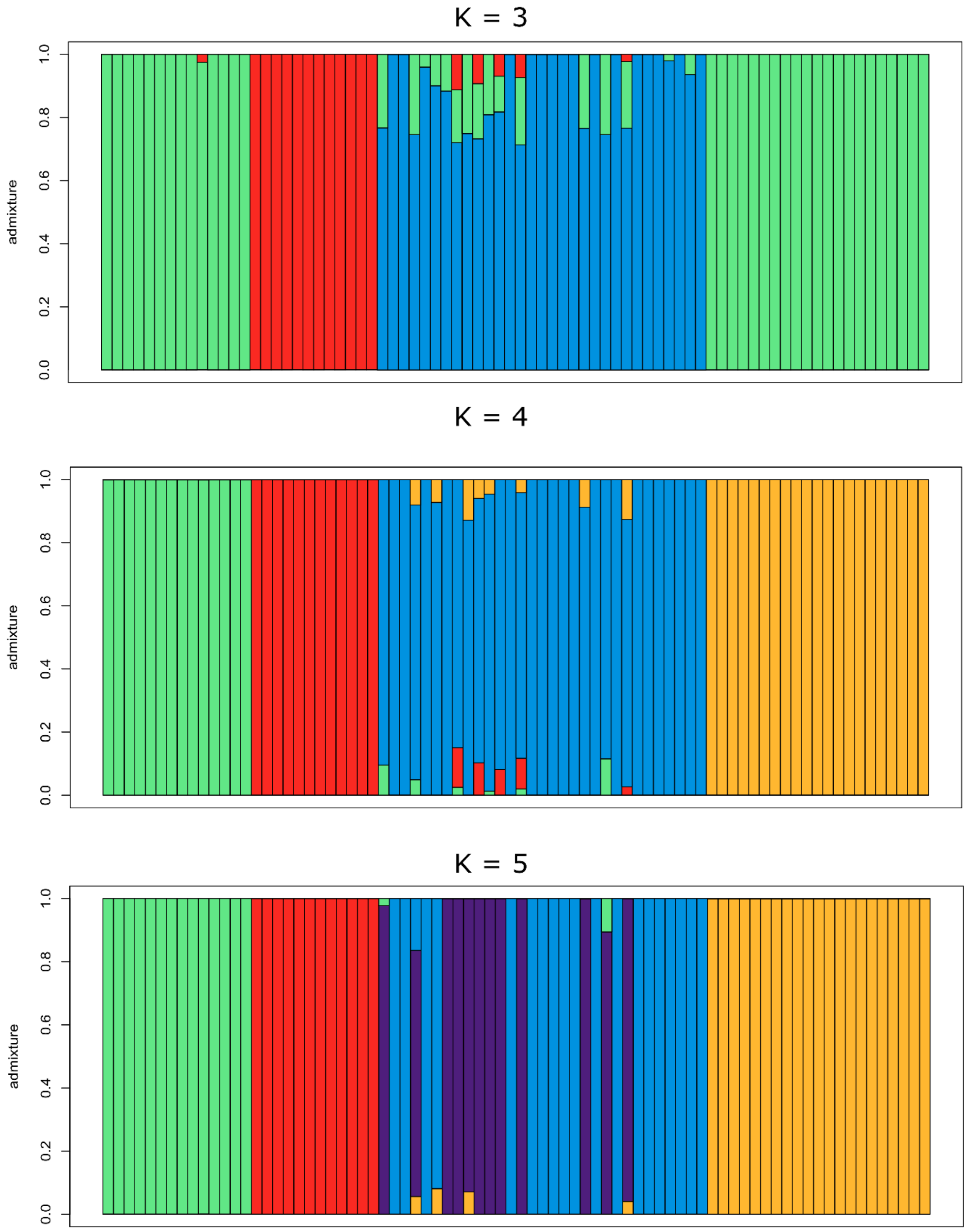


**Figure S12.** fastSTRUCTURE plot showing the best-fitting clustering models (K = 3, K = 4, and K = 5, respectively). Species are identified by color: Blue and purple: *A. laetissimus*, green: *A. arsyecue*, orange: *A. nahumae* and red: *A. carrikeri*.

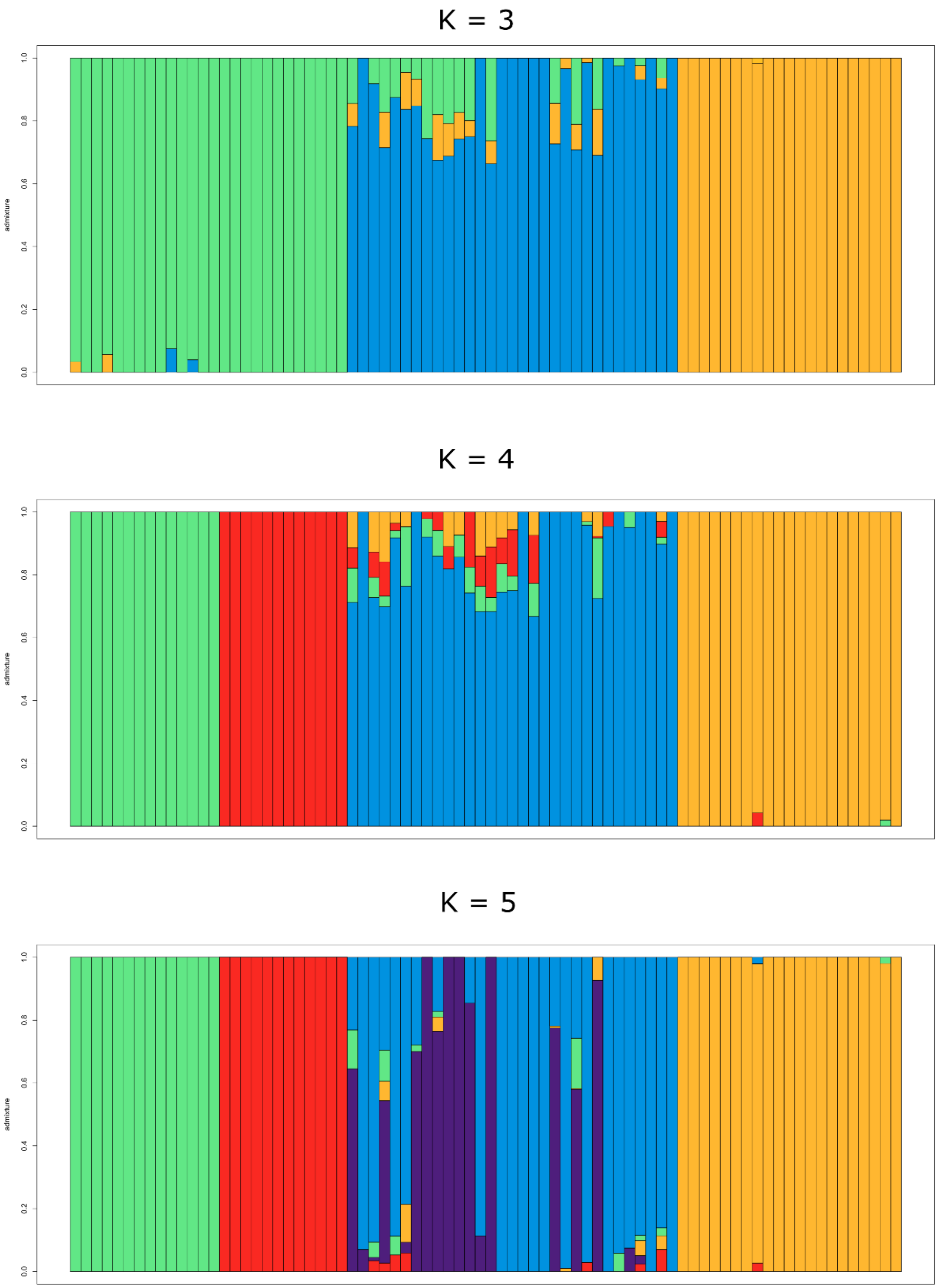


**Figure S13.** ADMIXTURE plot for the best-fitting clustering models (K = 3, K = 4, and K = 5, respectively). Species are identified by color: Blue and purple: *A. laetissimus*, green: *A. arsyecue*, orange: *A. nahumae* and red: *A. carrikeri*.

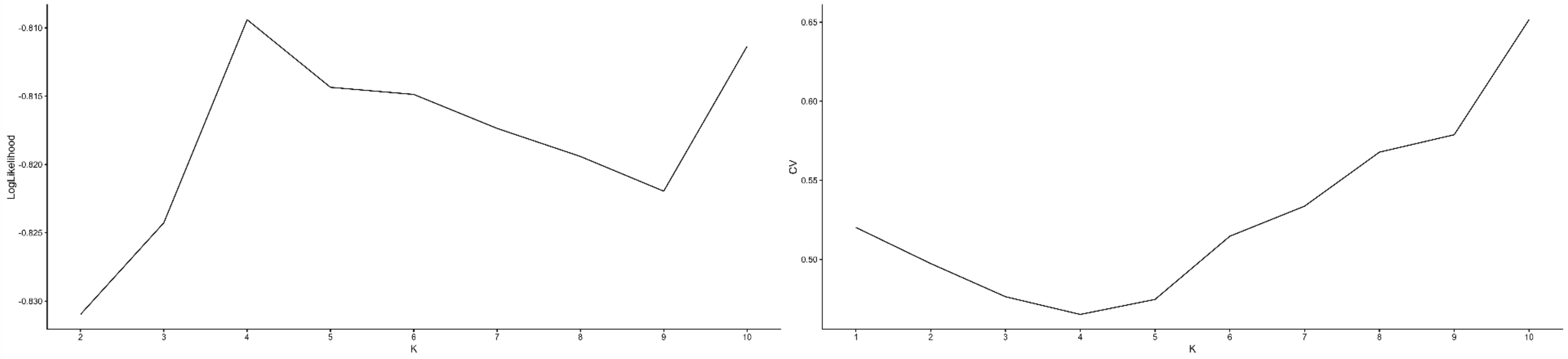


**Figure S14.** Determination of the most likely number of genetic clusters (K) using population structure analysis. (A) Log-likelihood values estimated by fastSTRUCTURE for K = 2–10. (B) Cross-validation (CV) values estimated by ADMIXTURE for K = 1–10. In fastSTRUCTURE, the maximum log-likelihood value was observed at K = 4, while in ADMIXTURE the lowest CV value was also recorded at K = 4.

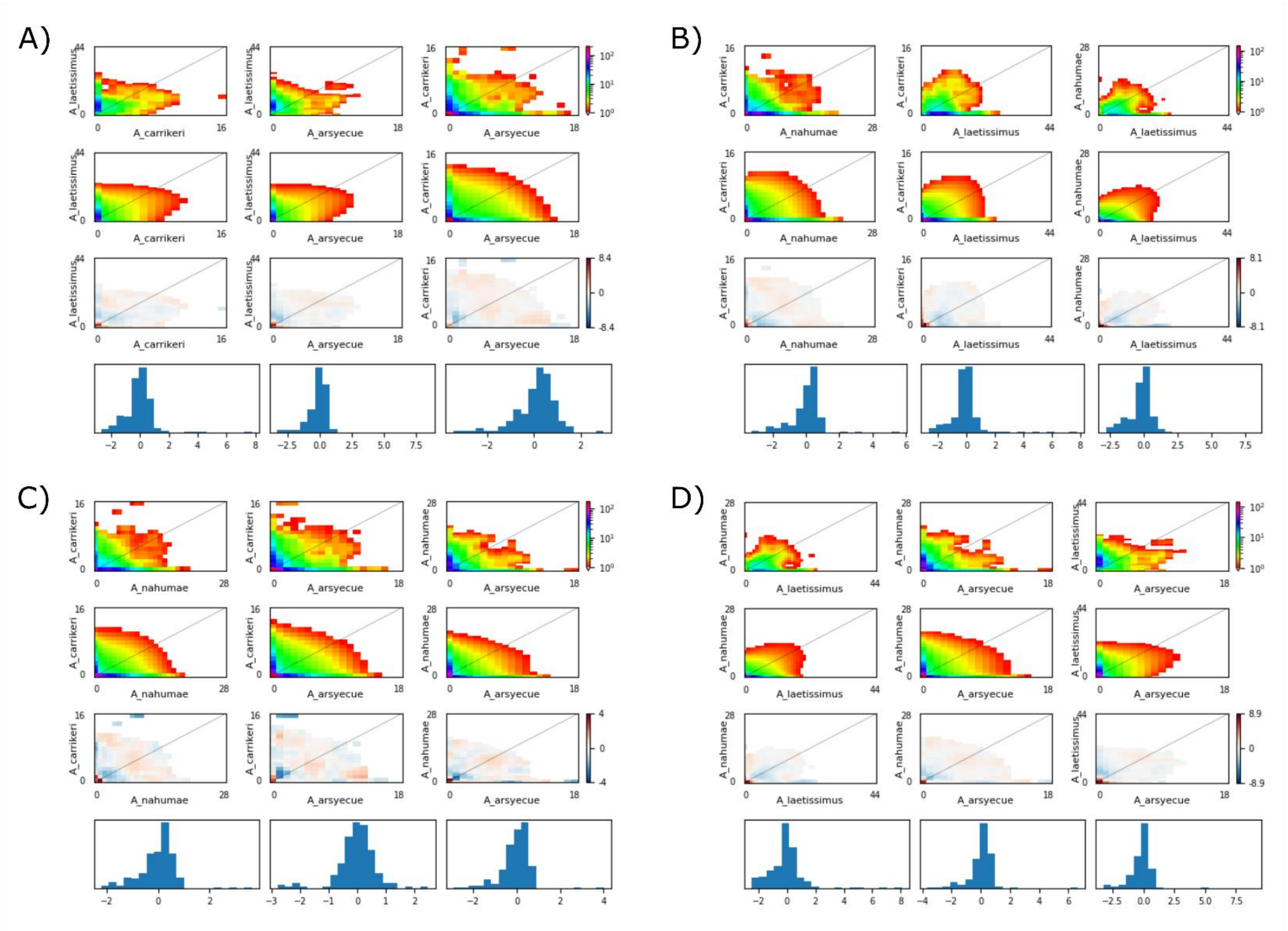


**Figure S15.** Observed (top row) and simulated (second row) joint site frequency spectra (JSFS) and residual plots (bottom rows) for each comparison in each of the four simulations conducted for three species.

**Table S3.** Estimated parameters from the best models for each of the four demographic history simulations conducted for three species, inferred using GADMA.

| Simulación-Morfoespecies | # modelos evaluados | Log-likelihood | AIC | Ne_ances | t_1 | Ne_1 | Ne_2 | Ne_3 | t_2 | m1_23 | m23_1 | m2_12 | m2_13 | m2_21 | m2_23 | m2_31 | m2_32 |
|---|---|---|---|---|---|---|---|---|---|---|---|---|---|---|---|---|---|
| *A. laetissimus - A. arsyecue - A. carrikeri* | 13274 | -2209.19 | 4466.38 | 24461 | 1410937 | 3264 | 12301 | 11343 | 47055 | 19.98 | 1999 | 0.94 | 0.95 | - | 1.70 | 3.32 | 1.34 |
| *A. laetissimus - A. carrikeri - A. nahumae* | 18570 | -2143.12 | 4334.24 | 45917 | 966896 | 6569 | 4039 | 11435 | 48553 | 19.97 | 2001 | 0.26 | 1.89 | 1.76 | 0.67 | 4.98 | 3.31 |
| *A. arsyecue - A. laetissimus - A. nahumae* | 15158 | -2214.55 | 4477.1 | 42244 | 977705 | 4270 | 9490 | 3518 | 132816 | 65.03 | 1992 | 1.46 | - | 4.47 | 3.29 | 1.66 | 0.75 |
| *A. laetissimus - A. carrikeri - A. nahumae* | 10074 | -1411.45 | 2866.89 | 31123 | 849712 | 3690 | 1247 | 2306 | 227239 | 203.24 | 2004 | 1.50 | - | 0.21 | 0.25 | 1.46 | 1.13 |

AIC: Akaike information criterion; Ne_ances: Ancestral population size before the first divergence event; t_1: Time since the first divergence event; Ne_1: Current effective population size of population 1; Ne_2: Current effective population size of population 2; Ne_3: Current effective population size of population 3; t_2: Time since the second divergence event; m1_23: Proportion of migrants per generation from ancestral population 1 to ancestral populations 2–3; m23_1: Proportion of migrants per generation from ancestral populations 2–3 to ancestral population 1; m2_12: Proportion of migrants per generation from population 1 to population 2; m2_13: Proportion of migrants per generation from population 1 to population 3; m2_21: Proportion of migrants per generation from population 2 to population 1; m2_23: Proportion of migrants per generation from population 2 to population 3; m2_31: Proportion of migrants per generation from population 3 to population 1; m2_32: Proportion of migrants per generation from population 3 to population 2. The proportion of migrants per generation was estimated as *4Nem*, where $N_e$ represents the effective size of the source population and *m* the migration rate to the receiving population.

**Table S4.** Estimation of ABBA and BABA proportions and D-statistics for all possible combinations of three species, with the species *A. spurrelli* and *A. fronterizo* combined as the outgroup (ancestral allele) (Dtrios function). Associated Z-scores and p-values are reported.

| Population 1 | Population 2 | Population 3 | Outgroup | Dstatistic | Z-score | p-value | f4-ratio | BBAA | ABBA | BABA |
|---|---|---|---|---|---|---|---|---|---|---|
| *A. arsyecue* | *A. laetissimus* | *A. carrikeri* | *A. spurrelli - A. fronterizo* | 0.0751966 | 1.58385 | 0.113229 | 0.161215 | 25.6688 | 22.6434 | 19.4761 |
| *A. arsyecue* | *A. nahumae* | *A. carrikeri* | *A. spurrelli - A. fronterizo* | 0.0108137 | 0.168055 | 0.86654 | 0.0216538 | 21.5439 | 19.8828 | 19.4573 |
| *A. arsyecue* | *A. laetissimus* | *A. nahumae* | *A. spurrelli - A. fronterizo* | 0.0458086 | 1.10822 | 0.267769 | 0.150673 | 24.3294 | 22.1651 | 20.2233 |
| *A. nahumae* | *A. laetissimus* | *A. carrikeri* | *A. spurrelli - A. fronterizo* | 0.0633013 | 1.57219 | 0.115905 | 0.14689 | 23.889 | 23.0279 | 20.2861 |

**Table S5.** Estimation of ABBA and BABA proportions and the D-statistic without an outgroup, using the four species present in the SNSM as an unrooted tree and an arbitrary assignment of the ancestral allele (Dquartets function). Associated Z-scores and p-values are reported.

| Population 1 | Population 2 | Population 3 | Population 4 | Dstatistic | Z-score | p-value | f4-ratio | BBAA | ABBA | BABA |
|---|---|---|---|---|---|---|---|---|---|---|
| *A. laetissimus* | *A. arsyecue* | *A. carrikeri* | *A. nahumae* | 0.0162429 | 0.673135 | 0.500861 | 0.0391 | 47.6300 | 47.6085 | 46.0866 |

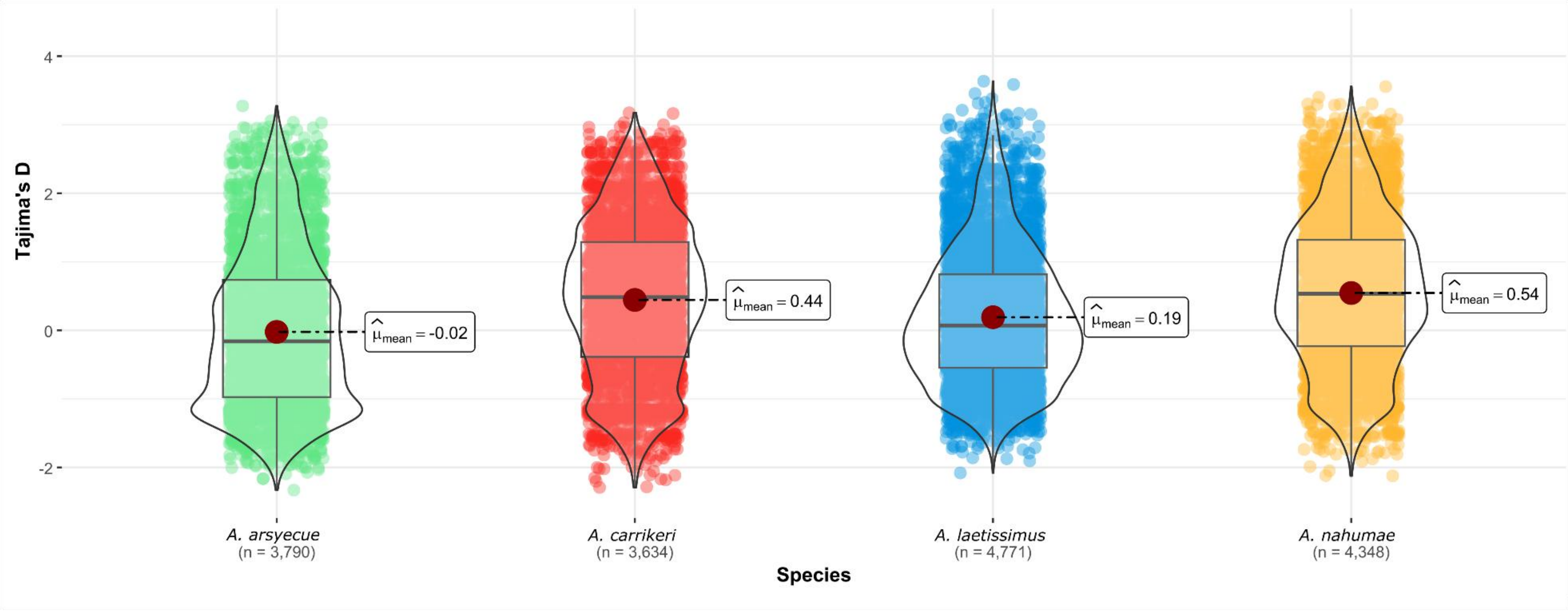


**Figure S16.** Tajima's D statistic by tag for each species. The red circles indicate the mean. Species are identified by color: Blue: *A. laetissimus*, green: *A. arsyecue*, orange: *A. nahumae* and red: *A. carrikeri*.

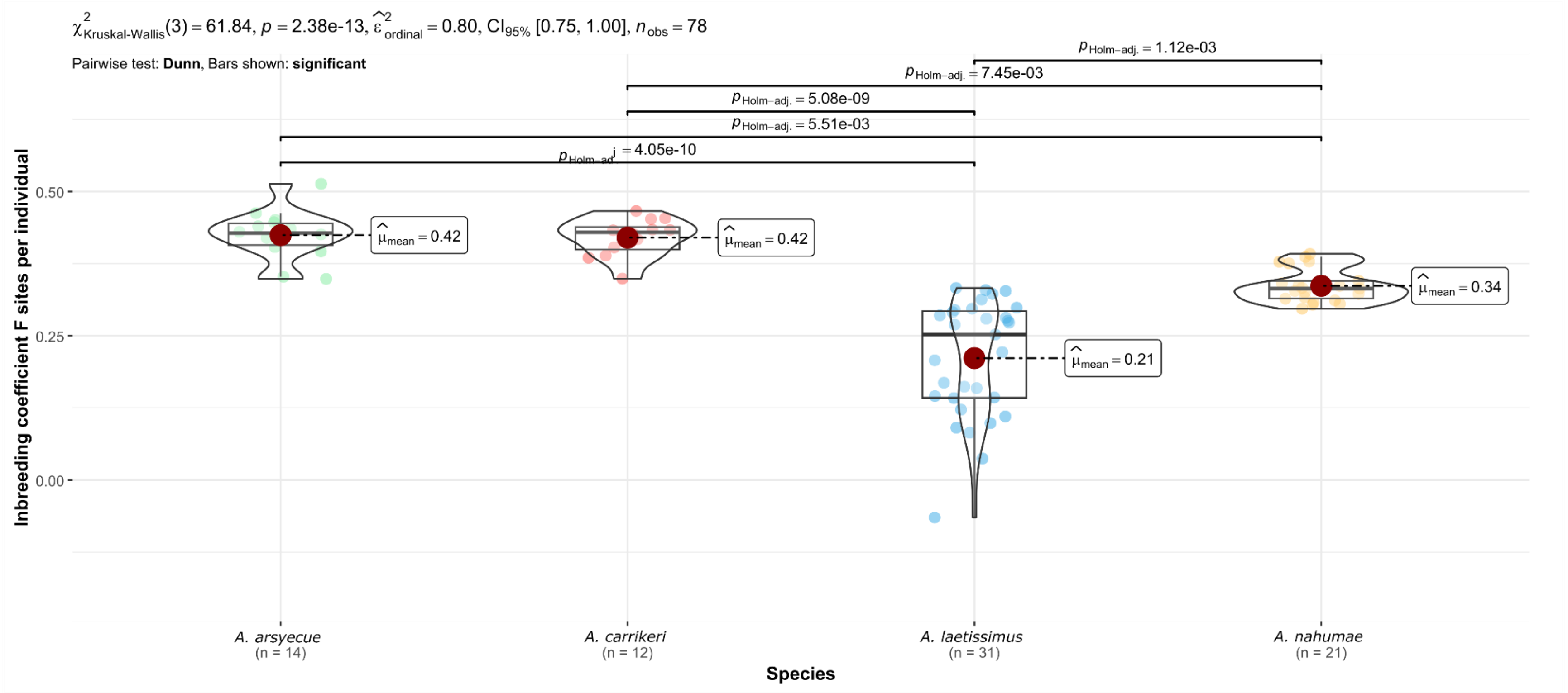


**Figure S17.** Inbreeding coefficient $F_{(HOM)}$ by individual per species. The red circles indicate the mean, the black horizontal line indicates the median, and the upper bars show the statistical significance of the differences. Species are identified by color: Blue: *A. laetissimus*, green: *A. arsyecue*, orange: *A. nahumae* and red: *A. carrikeri*.
